# Expansion, restructuring and characterization of the *Legionellaceae* family

**DOI:** 10.1101/2024.10.21.619444

**Authors:** Marco Gabrielli, Alessio Cavallaro, Frederik Hammes

## Abstract

*Legionellaceae* is a large and widespread family of facultative intracellular bacteria with high clinical relevance. While *Legionella pneumophila*, the most clinically relevant species, is relatively well studied, very limited information is available regarding the other species belonging to this family, despite its only recognized genus (i.e., *Legionella*) being included in water regulations worldwide. Here, we analyzed publicly-available *Legionellaceae* isolates and metagenome-assembled genomes, nearly doubling the number of recognized species and highlighting the need to consider restructuring the family’s taxonomy to include multiple genera that present distinct clinical relevance. We further found that, except for Type II and IVB secretion systems, traits linked to host invasion vary significantly across species, likely influencing their host range and survival in the environment. Presumptive genus clusters exhibited variations in metabolic capabilities, but *L. pneumophila* and closely related species generally displayed more complete pathways and fewer auxotrophies, suggesting a less strict intracellular lifestyle. On the contrary, specific deep-branching clades show reduced genome size and the lack of poly-3-hydroxybutyrate biosynthetic potential suggesting a stronger host association. Finally, we showed that most species are present across various environments and that, while they have apparently distinct environmental distributions, niche overlap increases at higher metabolic similarity. Together, our results shed light on the ecology of these microorganisms, highlight the diversity of traits that can occur in closely related facultative intracellular bacteria, and call for the restructuring of this family to align with genomic information and clinical relevance to aid the management of *Legionellaceae* bacteria.

**Importance:** Species belonging to the *Legionellaceae* family are responsible for Legionnaires’ disease, the leading waterborne disease in many industrialized countries. Historically, the focus was (and still is) on *Legionella pneumophila,* and the ecology of other species remained largely unknown and unexplored. However, several water safety regulations cover the entire *Legionella* genus,currently the only recognized within this large family, thus including species with unknown health relevance and ecology. This work provides an in-depth characterization of the genomic diversity across the family *Legionellaceae*, highlighting the concentration of clinically-relevant species in specific clades presenting distinct putative phenotypes linked to host invasion, metabolism and environmental niches. These findings highlight the need to consider restructuring the taxonomy of the family *Legionellaceae* in multiple genera to prioritize interventions to control the presence of the most clinically-relevant ones and advance public health protection.

## 1. Introduction

Microbial ecology largely relies on short-read 16S rRNA sequencing and specific model organisms to infer microorganisms’ environmental distribution and phenotypic characteristics. While this combination has enabled important ecological insights (1,2), the limits imposed by amplicon sequencing taxonomic resolution (3) and the restricted number of model organisms available (4) hamper our understanding of the ecological differences at lower taxonomic levels (5,6). When only some of the species within a clade are of interest (e.g., clinically relevant), the average response of the clade may contrast with the response of the individual species, as in Simpson’s paradox (7). As a consequence, the data obtained from studying the clade may not always properly inform legislation and management practices targeting individuals.

A relevant example of this problem lies in the family *Legionellaceae* and its only recognized genus, *Legionella*. Multiple *Legionella* species can cause legionellosis, a disease responsible for a significant number of deaths and costs exceeding 800 million USD per year in the USA alone (8) and for which the reported incidence is rising worldwide (9). While about 70 species are currently recognized (10), *L. pneumophila* is alone responsible for more than 90% of documented infections in Europe and the USA (11) and the remaining cases have been attributed to about 40 other species (e.g., about 50% of isolated species) (12). This discrepancy between the large number of species and the clinical importance of *L. pneumophila* results in different legislative actions, with countries differing with regards to the regulatory target (e.g., only *L. pneumophila*, *Legionella* spp., or a combination of the two) (13,14), as well as the regulated exposure source (e.g., showers, drinking water, cooling towers) (15). While a broader regulatory target (i.e., *Legionella* spp.) follows a precautionary principle, an overly broad target can conflate different species with distinct phenotypes and hamper prioritisation of truly critical situations. However, since information on non-*L. pneumophila* species is scarce, it is currently impossible to assess both their potential as “indicators” of *L. pneumophila*, as well as the efficacy of *L. pneumophila* control strategies on other pathogenic species. Only by obtaining a deeper understanding of the ecological differences among different *Legionellaceae* species it will be possible to fill such knowledge gaps and improve the management and legislation of these species (13).

So far, most of the information available on non-*L. pneumophila* species has been obtained from comparative genomic approaches on isolated species. These studies uncovered significant heterogeneity in terms of genome organization (16), encoded effectors (17) and secondary metabolites (18). However, information on the factors which affect their interactions with the environment and potential hosts, such as motility, secretion systems (beyond the conserved Type II and IVB (19,20)) and metabolism, is still lacking. Moreover, the aforementioned studies focus only on cultured isolates, while recent molecular-based surveys have uncovered a significant uncultured diversity of *Legionellaceae* bacteria (21).

Here we retrieved over 5000 publicly available *Legionellaceae* genomes and metagenome-assembled genomes and compared the functional potential of the different species included in this family, focusing specifically on evaluating its genetic diversity and different traits important for facultative intracellular bacteria, including secretion systems, motility and host invasion, and metabolism. We complemented this analysis with a genome-informed mining of 16S rRNA surveys, assessing species’ distribution across environments. Collectively, this work provides key insights regarding the diversity and the ecology of *Legionellaceae* bacteria which will boost our knowledge regarding the environmental lifestyles of *Legionellaceae* bacteria and allow to refine legislation and management strategies towards the most clinically relevant species to safeguard public health.

## 2. Results and discussion

### 2.1 Analysis of public genomes substantially restructures the *Legionellaceae* family

Given the known difficulties in culturing bacteria (22), culture-independent data is necessary to characterize the full range of microbial diversity. We delineated 129 species clusters (Table S1) from publicly available *Legionellaceae* genomes, both isolated and metagenomic-assembled (MAGs), using an average nucleotide identity (ANI) of 95% (23). Importantly, ANI-delineated species clusters showed high concordance with existing taxonomic classifications, including genomes from a single species and/or without species-level annotations. The newly delineated species clusters substantially expand the *Legionellaceae* family, nearly doubling the number of the species currently recognized (24,25). Because many of these novel species form clades phylogenetically distant from currently isolated taxa, their inclusion expands the phylogenetic diversity of the family by 142% (Figure 1). At the same time, we identified several new species clusters phylogenetically related to known ones. As an example, ANI similarity supports the differentiation of *L. pneumophila* subspecies *subsp. pascullei* and *subsp. fraseri* into two distinct species (i.e., *L. pascullei* and *L. fraseri*) based on an average ANI similarity to *L. pneumophila* of, respectively, 91.2% and 92% (i.e., below the accepted 95% ANI threshold). Notably, given that most of the unique genes are of unknown function (26), the phenotypic differences linked with this differentiation are unknown. Importantly, while uncultivated *Legionellaceae* species increase the diversity of the family, they do not impact family-level boundaries with sister clades: all *Legionellaceae* genomes were separated from the outgroup genomes derived from the genera *Coxiella*, *Berkiella* and *Aquicella* with high confidence (bootstrap = 100).

**Figure 1.**
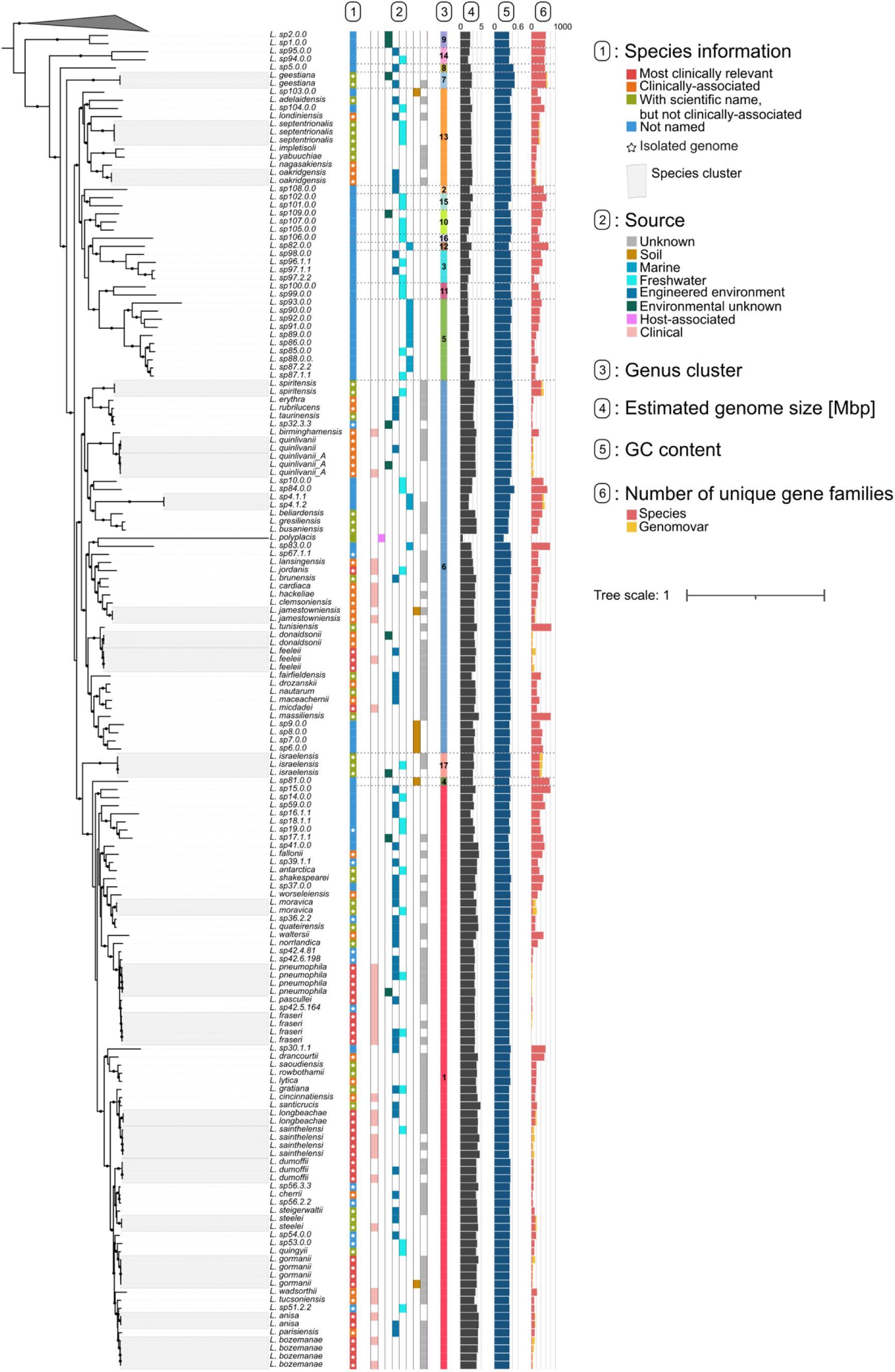
Clinical relevancy, genome origin, genus clusters and genome characteristics of the *Legionellaceae* genomes analyzed. Maximum likelihood phylogenetic tree based on 369 BUSCO *Legionellales* protein families (123421 sites). Branches with optimized ultrafast support ≥95% are marked with black circles. Leaves enclosed within grey areas indicate genomes of different genomovars belonging to the same species. Outer tree layers indicate whether the delineated species have currently valid scientific names and genomes from isolated specimens are available, the source of all the genomes belonging to the specific species/genomovar, genus cluster membership (cluster number highlighted once per clade), the estimated genome size and GC content, and the number of unique gene families present throughout a species or unique to specific genomovars. The collapsed clade represents the outgroup, while horizontal dotted lines highlight the genus cluster boundaries.

Recognizing the lack of a consensus definition, a genus is generically considered a restricted monophylic clade of organisms with conserved characteristics, which arise due to significant genome similarity (27). The family *Legionellaceae* currently includes only a single genus, i.e. *Legionella* (24,25). However, a comparison of the average amino acid identity (AAI) returned values reaching as low as 45%, a value well below the suggested genus boundary (i.e, 65 - 72%) (28). Similar evidence was provided by the percentage of conserved proteins (POCP) down to 18%, much lower than the accepted genus-wide threshold of 50% (29). We then clustered genomes into genus clusters based on a 50% POCP threshold, obtaining clusters which, except for rare cases, followed the accepted 50% genus threshold (Figure S1). This resulted in 17 presumptive genus clusters identified within the *Legionellaceae* family, of which only five include cultured isolates. This is substantially different from what is currently accepted, and highlights the need to consider restructuring the taxonomy of this family. This restructuring is concordant with previous efforts in bacterial taxonomy and *Legionellaceae* comparative genomics, underlining the robustness of our results (Supplementary Text 1). While clinical relevancy was not considered during genus clusters definition, most clinically-relevant species clusters (30) belong to the same clade as *L. pneumophila.* However, clinically-associated species are present in three of the five genus clusters with isolated representatives. Noteworthy exceptions are *L. micdadei* and *L. feeleii* which were found to belong to a separate genus cluster than *L. pneumophila,* despite being two of the most clinically relevant species after *L. pneumophila* (34,35). The absence of clinical isolates from a large clade of the *Legionellaceae* tree suggests that the current definition of the *Legionella* genus (i.e., coinciding to the family one) includes clades with a clear distinction in clinical relevance. Consequently, in case legislation is applied at the genus level, the risk of *L. pneumophila* and species with no indication of clinical relevance is equated, not allowing legislators/operators to prioritize remediation interventions.

We found the genus clusters to present distinct genome characteristics, indicating different evolutionary pressures likely due to distinct lifestyles (33). Genome size and GC content varied significantly across genus clusters, even when accounting for genome origin (i.e., isolates, MAGs) (two-way ANOVA, p-vals < 0.001) (Figure 1). The genus clusters of *L. pneumophila* presents significantly larger genome sizes and lower GC content than other genus clusters including both previously isolated species (i.e., *L. micdadei* and *L. oakridgensis*), as well as novel ones (genus clusters 6, 13, 5; Tukey HSD test, p-val < 0.003). The proteins encoded by different species were also found to be very diverse: out of 37447 protein families only 843 (2.2%) core proteins are conserved (i.e., present in ≥95% of genomes). To put this number into context, a recent analysis showed the family *Coxiellaceae* to present 348 core protein families (34), a comparable number when accounting for the different number of genomes analyzed across clades (333 predicted core proteins in *Legionellaceae*). This similarity is consistent with the high phenotypic diversity found in both clades. On the other hand, a similar percentages of conserved proteins has been found only in the phylum *Chlamydiae* (35), a much broader taxonomic clade, indicating the considerable diversity of *Legionellaceae* species.

Comparative genomic efforts have been focused, so far, only on isolated *Legionellaceae* species (16,17,19,20). However, our data shows that this fails to capture a significant fraction of genomic diversity. In fact, the number of core and total protein families in the *Legionellaceae* family are, respectively, −26.1% and +30.4% different than what estimated based only on isolated species (core families: 1132, total families: 23021) (36). A smaller core and a broader diversity of protein families indicate that uncultivated species lack protein families conserved in their cultivated counterparts, while, at the same time, encode for distinct proteins potentially responsible for novel traits. The protein families from uncultivated species expand the known capabilities of *Legionellaceae* bacteria in virtually all aspects of cellular lifestyle as shown by the wide distribution of their functional categories (i.e., COG classes) (Figure S2). This diversity is mainly driven by shared phylogenetic history. The number of unique protein family per species (range: 2 – 637) is largely explained by the phylogenetic distance to the closest other *Legionellaceae* genome (R^2^ = 0.69, Figure S3). Given the high phylogenetic relatedness of the L. pneumophila genomes analyzed, this result explains the low number of unique proteins detected in those genomes (Figure 1). While genome size also influences the number of unique genes present, its influence is secondary (ΔAIC: genome size = 16.6; phylogenetic distance = 181.2). This result suggests that the increase in genome size was accomplished in *Legionellaceae* bacteria through different evolutionary trajectories which led to a diversification of proteome content.

### 2.2 Diversity in the *Legionellaceae* family highlights differences in putative phenotypes

The extent of the proteome diversity across different genus clusters suggests the presence of distinct phenotypes across this family. Given the importance of interactions between facultative intracellular bacteria with their host, as well as the importance of metabolism for bacterial niche definition, we assessed the differences of (i) secretion systems, (ii) factors linked to host invasion and (iii) the metabolic potential across different species and genus clusters.

#### 2.2.1 Secretion systems presence is highly variable within the *Legionellaceae* family

The ability of secretion systems to translocate effector proteins to host cells is a key trait of intracellular bacteria, which allows them to manipulate hosts to their advantage (37). *L. pneumophila* is known to secrete a variety of effectors through the Dot/Icm Type IVB (T4BSS) and Type II (T2SS) secretion systems (17,20,38). We found homologs of proteins belonging to the T4BSS of *L. pneumophila* in all genomes except *L. polyplacis*, an obligate insect symbiont (39), indicating that, despite being conserved throughout most of the family, this trait can be lost during the adaptation to specific intracellular niches. Besides the T4BSS secretion system, the T2SS was detected in nearly all *Legionellaceae* species, expanding previous knowledge about its distribution to most uncultured species (40). This secretion system was not detected in *L. polyplacis* and genus cluster 9. While the absence of T2SS in this uncultivated clade could be due to genome incompleteness (estimates: 11% and 1%), it is possible that such machinery was lost by the clade’s ancestor. In fact, T2SS is not strictly necessary for host entry above 22 °C, but rather favors *L. pneumophila* survival at cold temperatures (41). It is then possible that species lacking a T2SS might not be able to survive at low temperatures or are on an evolutionary path towards a stricter intracellular lifestyle. Interestingly, two of these MAGs were retrieved from environments with warm temperatures (42,43), supporting the role of T2SS in cold-adaptation.

Despite the potential role of other secretion systems in *Legionellaceae* ecology (e.g., host range, motility, HGT), little is known regarding their prevalence and diversity (44). Our results highlight the presence of a wide range of secretion systems throughout the family (Figure 2, Table S2), with different secretion systems having distinct distributions across genus clusters. We detected Type IVA secretion systems (T4ASS) of various types (i.e., Types F, G, I and T) throughout the family and found only a limited influence of phylogenetic relatedness on their presence/absence pattern (Jaccard diversity-based Mantel correlation, r = 0.18, p-val < 0.001) due to their frequent presence on plasmids (i.e., 78% of Type F and 43% of Type T T4ASS) (17,45). Conversely, we found other secretion systems to be present predominantly in specific genus clusters. We found the lss and novel Type I (T1SS) and Type V (T5SS) secretion system predominantly in the genus clusters including *L. pneumophila*, *L. micdadei* and *L. israelensis* (genus clusters 1, 6 and 17, respectively). Such machineries could be potentially favor host cell adhesion and entry (46), as well as biofilm formation (47). In addition, we identified Type VI secretion systems (T6SS) within restricted clades of genus clusters 1 and 6. T6SSs could potentially be used against either competing prokaryotes or non-infectable eukaryotes (48) or to obtain nutrients from neighboring cells (49), potentially favoring extracellular survival. On the other hand, we noticed different types of T1SSs (i.e., lb1ss, lb2ss and lrpss) (50) to be largely lacking in the *L. pneumophila* genus cluster, while being present in the ones of *L. micdadei*, *L. oakridgensis* and novel clades (genus clusters 6, 13, 3, 11, 12, 16).

**Figure 2.**
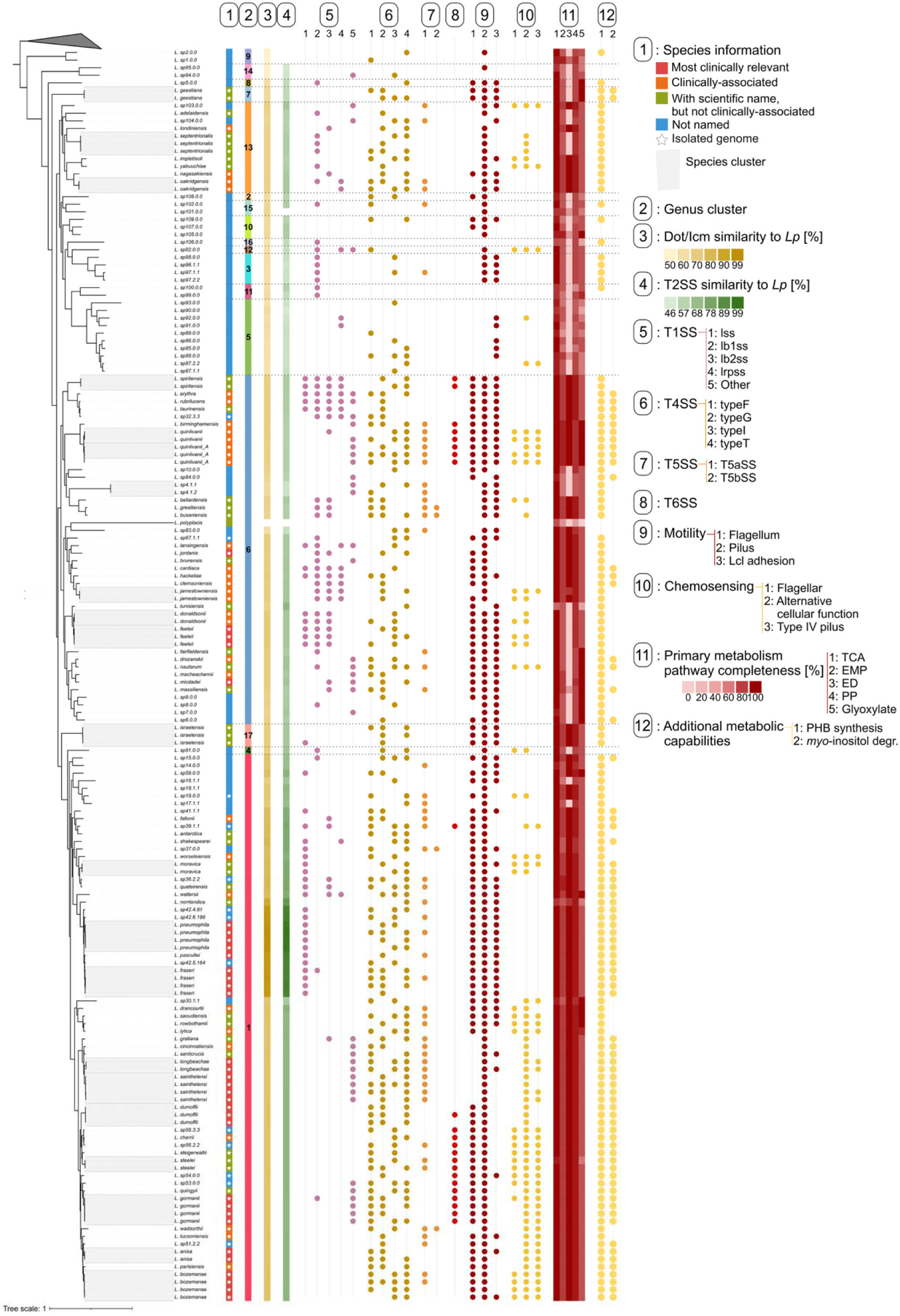
Distribution of inferred traits related to secretion systems, motility-related and metabolic among the analyzed *Legionellaceae* genomes. Maximum likelihood phylogenetic tree based on 369 BUSCO *Legionellales* protein families (123421 sites). Leaves enclosed within grey areas indicate genomes of different genomovars belonging to the same species. The average Dot/Icm and T2SS protein similarity was estimated against *L. pneumophila* strain *Paris*, while rounded markers indicate the presence of the specified traits.

A direct connection between the reduced presence of secretion systems in given clades and their pathogenicity cannot be drawn (51,52). Intracellular replication is connected to the release of adequate effectors within a host (53). Given the known heterogeneity of effectors even within single species (e.g., 20,32), encoding genes for a secretion system is not a sufficient proxy for released effectors.

#### 2.2.2 Factors linked to host invasion are not conserved among species

The likelihood of an intracellular bacterium encountering its host can be very low (55)(56), suggesting the importance of “active” host invasion mechanisms such as motility. We identified the genes required for Type IV pili in throughout the family except for a deep-branching clade composed of two novel genus clusters (5 and 11) (Figure 2, Table S2), as well as *L. polyplacis*, indicating widespread ability for twitching motility across the family. Conversely, we found flagellar assembly genes to be present mostly in the genus clusters of *L. pneumophila*, *L. micdadei*, *L. geestiana* and *L. israelensis* (genus clusters 1, 7 and 17) and 84% of the clinically-associated species. Flagellar assembly proteins are mostly absent in other genus clusters. Among the most clinically-relevant species, only *L. longbeacheae* and *L. sainthelensis* lack flagellar proteins. Notably, 62.5% of soil-isolated species, were found to be non-flagellated, suggesting flagella-independent surface motility as an adaptation to terrestrial environments (55). Beyond motility, adhesins, such as the Lcl protein (57), are also key determinants of host-pathogen interactions and biofilm formation (58). We found Lcl homologs characterized by a collagen-like domain only in a subclade of the genus cluster of *L. pneumophila*, while we detected additional novel proteins with collagen domains in 38 species across the family (Table S3). Enhanced motility and biofilm formation abilities provided by these proteins is likely to favor host colonization and resistance to stresses, leading, thus, to persistence in the environment. Yet, characterizing the importance of such traits is complicated by hosts diversity. For example, *L. pneumophila* was shown to require a flagellum to efficiently infect human macrophages, but not amoebae (59). However, the existence of clinical non-flagellated species indicates that flagella are not a strict requirement for human infectivity.

Chemosensing can be used to direct movements (i.e., chemotaxis) towards hosts (55). However, except for subclades within genus clusters 1 and 6, most *Legionellaceae* species lacked both CheW and methyl-accepting chemosensing proteins, two of the essential components of the chemotaxis Che system (60) (Table S3). In addition, we found weak associations between the Che pathway and the presence of flagella (binomial GLM, p-value = 0.02) and pili (p-value = 0.07), further stressing the unrelatedness of these traits. While chemotaxis might not be a requirement for pathogenic bacteria once in the respiratory system (57,61), this ability is likely key in supporting the survival in the environment of these bacteria (61). In fact, other regulatory pathways for chemotaxis might be present in *Legionellaceae* bacteria. For example, twitching motility in *L. pneumophila* is inhibited by the presence of other bacteria likely due to lower nutrients concentrations (62), indicating either that rich nutrient conditions are needed for twitching motility or the ability to perform chemotaxis despite lacking the Che pathway.

Beyond chemotaxis, we predicted up to 80 signal transduction and 170 response regulator proteins per genome, with *L. polyplacis* encoding for no signal transduction proteins and only five response regulators. We found a different number of these proteins across the isolated bacteria in different genus clusters (ANOVA, Tukey HSD post-hoc test, p-values < 0.002; Figure S4), with genus cluster 1 presenting the highest number. In total, we identified 255 signal transduction and 561 gene families for response regulation, of which only 3 and 12 were, respectively, detected across the whole family (≥95% species) and, In most cases, already reported for *L. pneumophila* (Table S4). Beyond those, we detected almost half of signal transduction (48%) and response regulators (42.4%) proteins to be genus cluster-specific, but that only respectively 40.3% and 24.4% of them to be encoded by multiple species within their genus cluster. These results suggest that while a restricted “core” regulatory network is conserved across species, significant diversity is present not only across, but also within genus clusters, which might significantly affect phenotypic responses to environmental cues and contribute defining the environmental niche of different species.

#### 2.2.3 Genus clusters exhibit metabolic potentials concordant with different degrees of host association

While *L. pneumophila* exhibits a complex biphasic metabolism (63), the metabolism of other species has been seldomly examined. We found that the tricarboxylic acid (TCA) cycle is complete in nearly all species, highlighting a family-wide reliance on serine for growth (64) (Figure 2, Table S5). Similarly, we found vitamin B biosynthesis pathways to be broadly conserved (Figure S5), including in the obligate symbiont *L. polyplacis* (39). Incomplete pathways were observed only in specific genus clusters (i.e., 3, 9, 14) or genomes (Figure S5), though further study is needed to elucidate whether these reflect host adaptation or environmental vitamin uptake.

Other metabolic pathways were far less conserved across the family. The Entner-Doudoroff (ED), Embden-Meyerhof-Parnas (EMP) and pentose phosphate (PP) pathways are used by *L. pneumophila* during post-exponential phase to switch from a replicative to an infectious phase (71,70,72). Yet, only highly-related species to *L. pneumophila* are likely to adapt the same strategy to switch life phases given the negative correlation between their completeness and phylogenetic distance to *L. pneumophila* (Spearman correlation; ρ = −0.48 – −0.7, p-vals < 0.001). The completeness of these pathways varies significantly across genus clusters (Wilcox test; p-vals < 0.001). For example, complete ED pathways are restricted only to subclades of the genus clusters 1, 6, 13 and 17, and completely absent from the ones composed only of novel species (genus clusters 2 - 5, 10 - 12, 15, 16) signaling differences in their involving glucose metabolism (63). These results indicate that species belonging to other genus clusters rely on different metabolic pathways to switch life phase. For example, while the glyoxylate cycle is absent in *L. pneumophila* (64), we observed its complete set of genes in several species belonging mostly to genus clusters 6 - 8. This pathway could potentially contribute to satisfy carbon requirements in the absence of glucose.

Metabolic diversity across *Legionellaceae* genus clusters extends beyond these central pathways and involve other phenotypes reported for *L. pneumophila*. For example, the ability to synthetize PHB and metabolize *myo*-inositol, key factors for intracellular replication and survival in nutrient-poor conditions (80,77), tends to be rarer in genus clusters phylogenetically distant from genus cluster 1 (Figure 2), suggesting further metabolic differences across genus clusters. In fact, even when excluding MAGs to avoid incompleteness bias (69), we observed significant differences in the completeness of KEGG metabolic pathways across genus clusters (Figure 3a; PERMANOVA, R^2^ = 24.6%, p-val < 0.001) due to several cluster-specific enzymes (Figure S6). The genus clusters including *L. pneumophila* and *L. micdadei* (genus clusters 1 and 6) present greater metabolic capabilities compared to other clusters (Table S6). In particular, we found more complete pathways for phosphatidylcholine biosynthesis and acylglycerol degradation in genus cluster 1, while a higher completeness of the glycogen biosynthesis pathway in cluster 6, factors related, respectively, to virulence and vacuole biogenesis (70,71), as well as survival under nutrient starvation (72). Notably, metabolic similarity did not align with clinical relevance, as subclusters with comparable metabolic profiles spanned species with distinct clinical associations (Figure S7, Table S7), suggesting metabolic potential influences niche overlap rather than pathogenicity.

**Figure 3.**
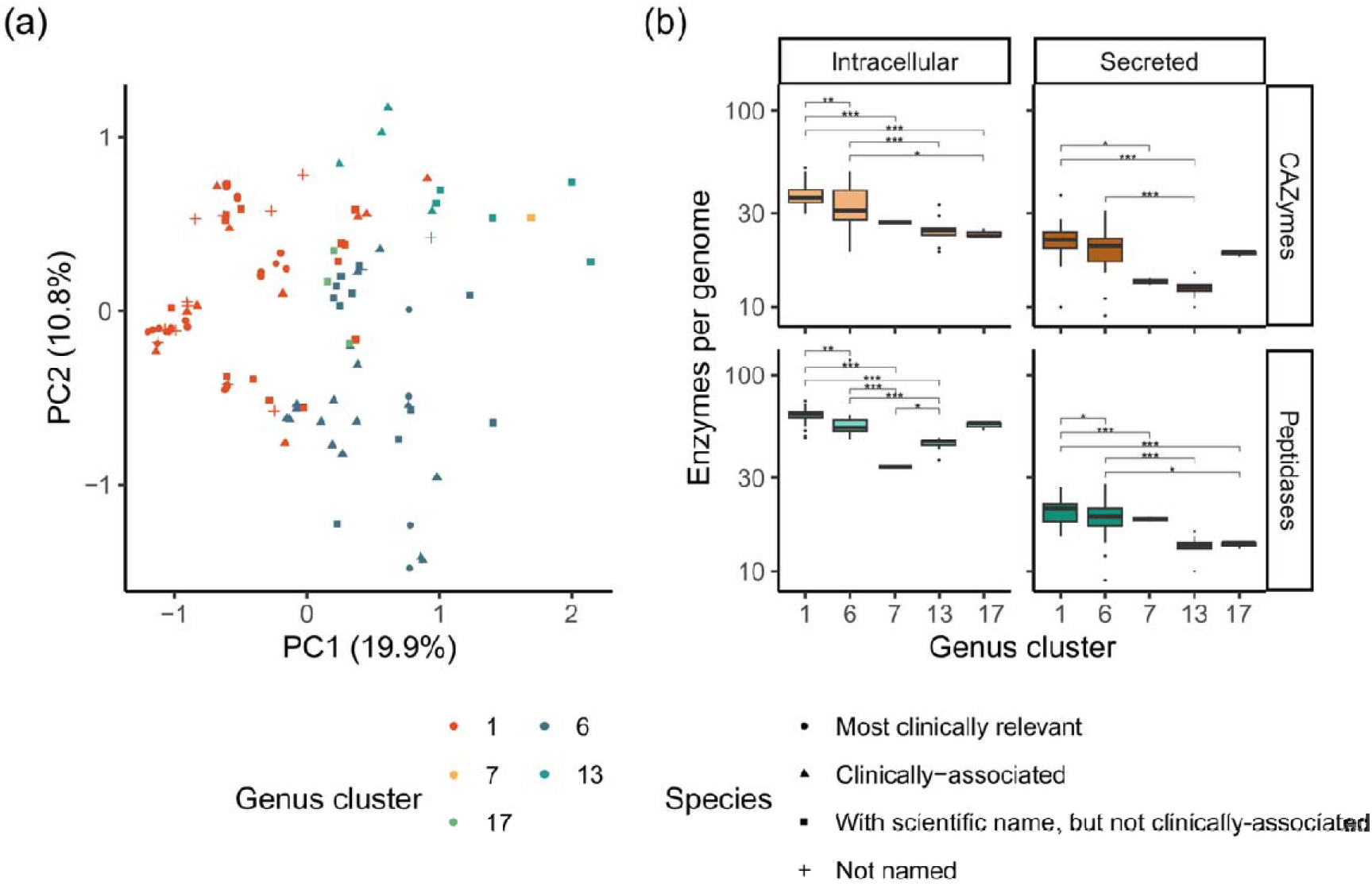
(a) Metabolic differences among *Legionellaceae* isolated genomes and clustering following genus cluster membership. (b) Number of CAZYmes and peptidades per isolated genomes as a function of their secretion and genus cluster membership. Statistically significant differences among genus clusters are marked by their p-values (Dunn test): ”*” = 0.05 - 0.01, ”**” = 0.01 - 0.001, ”***” < 0.001. Genus clusters representatives are: “1”: *L. pneumophila*, “6”: *L. micdadei*, “7”: *L. geestiana*, “13”: *L. oakridgensis*, “17”: *L. israelensis*.

The expanded metabolic capabilities of the *L. pneumophila* and *L. micdadei* genus clusters (genus clusters 1 and 6) include enhanced carbohydrates and peptide degradation, as shown by the significantly higher number of secreted and intracellular CAZymes and peptidases per genome (Wilcox test; p-vals < 0.001, Figure 3b). A greater number of CAZymes and peptidases, as well as their greater diversity (Figure S8) highlights the different abilities to degrade complex organic matter by different clusters (73,74).

Overall, the more restricted metabolic potential of deep-branching genus clusters and the lack of biosynthetic potential of energy storage compounds such as PHB, coupled with a reduced genome size, suggests their higher metabolic dependence on other microorganisms and their poorer survival in environments characterized by low nutrients concentrations and/or other stresses (75). While their sustenance through metabolites leaked from other microorganisms cannot be completely rules out (76), this might indicate a stricter intracellular nature and explain the difficulties in their isolation.

#### 2.2.4 Novel genus clusters show more auxotrophies than the ones including cultivated species

Intracellular bacteria have evolved auxotrophy for diverse amino acids (77), gaining a fitness advantage in their host-associated niche, but becoming dependent on external resources for survival (78). This also holds true for *L. pneumophila*, for which several studies have confirmed its auxotrophy for several amino acids (Table 1). We performed *in silico* analysis of *L. pneumophila* auxotrophies using GapMind, a pathway annotation tool (79), and gapseq, an automated tool for genome-scale metabolic model reconstruction (80). Except for L-isoleucine and L-serine, the predictions of the two tools were mostly concordant and matched, respectively, eight and six of the 11 reported auxotrophies, possibly due to strain-level differences (Table S8), as also highlighted by the inconsistencies among experimental studies. Discrepancies for L-isoleucine and L-serine arise because gapseq predicts two pathways not considered by GapMind (Table S10) which should warrant further biological validation. Interestingly, while L-cysteine is commonly used for culture confirmation of *L. pneumophila*, both tools did not predict *L. pneumophila* auxotrophy towards this amino acid. We detected the genes required for L-cysteine biosynthesis through the reverse transsulfuration pathway (81) in all *L. pneumophila* genomes. However, these genomes lack the capability to synthetize homocysteine, a required precursor for L-cysteine biosynthesis, as well as relying on inorganic sulphur sources (e.g., SO ^2-^, H S) (82), an aspect not controlled for in growth media recipes (e.g., BYEB). Such results indicate that L-cysteine might be an “indirect” auxotrophy resulting from the lack of inability to synthetize homocysteine and appropriate sulfur sources.

**Table 1.** Reported and predicted auxotrophies for *L. pneumophila*. Experimental evidence retrieved from experimental studies (64,83,84). High and medium GapMind biosynthesis prediction confidence results indicate the detection of homologs for all biosynthetic steps for a given amino acid. Conversely, low confidence results highlight the lack of biosynthetic steps and imply auxotrophy(79). Multiple results per amino acid indicate that distinct results were obtained with different genomovars. The asterisk indicates that while the genes for the L-cysteine biosynthesis have been annotated, the genes required to produce the required compound homocysteine are absent.

| Amino acid | Experimental evidence | GapMind biosynthesis prediction confidence | gapseq prediction |
| --- | --- | --- | --- |
| L-alanine | - | Not assessed | - |
| L-arginine | Auxotrophy consistently reported | Low | Auxotrophy |
| L-asparagine | - | High | - |
| L-aspartate | - | Not assessed | - |
| L-cysteine | Auxotrophy reported | Medium* | - |
| L-glutamate | - | Not assessed | - |
| L-glutamine | - | High | - |
| Glycine | - | High | - |
| L-histidine | - | Medium/High | - |
| L-isoleucine | Auxotrophy consistently reported | Low | - |
| L-leucine | Auxotrophy consistently reported | Low | Auxotrophy |
| L-lysine | - | High | - |
| L-methionine | Auxotrophy consistently reported | Low | Auxotrophy |
| L-phenylalanine | Auxotrophy reported | Low | Auxotrophy |
| L-proline | Auxotrophy reported | High | - |
| L-serine | Auxotrophy reported | Low | - |
| L-threonine | Auxotrophy reported | Low | Auxotrophy |
| L-tryptophan | - | Medium | - |
| L-tyrosine | Auxotrophy reported | High | - |
| L-valine | Auxotrophy consistently reported | Low | Auxotrophy |

Given the susceptibility of annotation-based tools to MAGs incompleteness and the tools’ overall concordance (Table S8), we continued the analysis only using gapseq’s results. The gapfilling carried out by this tool, in fact, removed the differences in the number of predicted auxotrophies between isolates and MAGs from the same genus cluster (t-test, p-vals > 0.1). This allowed us to highlight significant differences in the number of auxotrophies among genus clusters: we detected a lower amount in the genus clusters of *L. pneumophila*, *L. micdadei* and *L. israelensis* (genus clusters 1, 6, 17) compared to the ones composed exclusively of novel species (ANOVA, p-val < 0.001). We did not find any amino acid auxotrophy to be conserved across the entire family, despite several being highly prevalent (Figure 4, Table S10). Conversely, no species, including the obligate endosymbiont *L. polyplacis*, showed auxotrophy towards L-alanine, L-aspartate, L-glutamate, L-glutamine and glycine. Similarly to what reported for *L. pneumophila* (see above), we did not detect L-cysteine auxotrophy in genus cluster 1 indicating that such an “indirect” mechanism of L-cysteine auxotrophy extends also to other related species. However, we found “direct” L-cysteine auxotrophies in the genus cluster of *L. micdadei* and in several others without isolated representatives. Excepting cluster 7 (*L. geestiana*), cultured genus clusters rarely exhibited L-histidine auxotrophy, whereas uncultured clusters frequently displayed L-histidine and additional auxotrophies. The additional auxotrophies in yet-to-be-cultured genus clusters calls the supplementation of such media with additional amino acids for the isolation of novel species.

**Figure 4.**
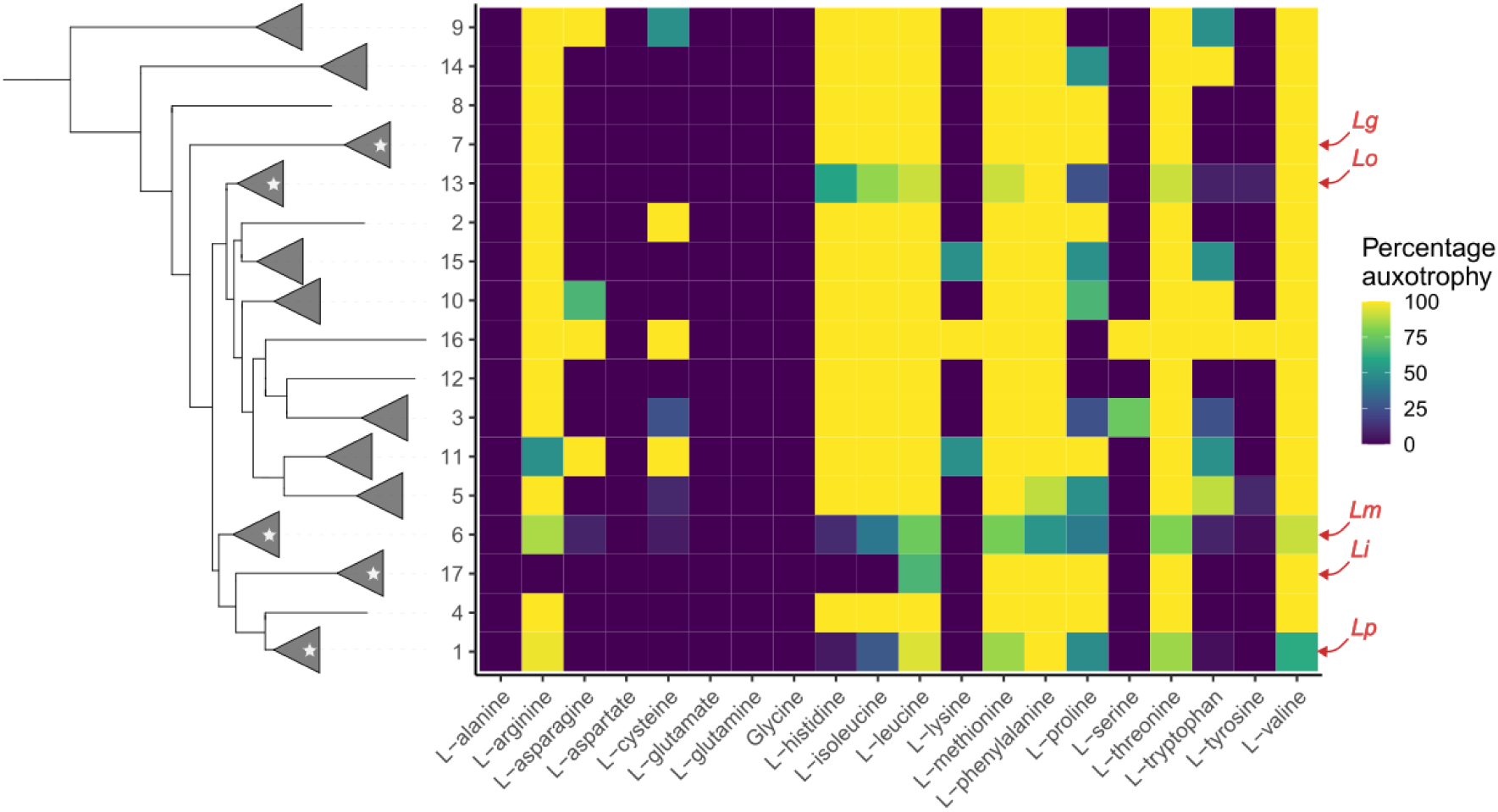
Prevalence of auxotrophies predicted in each genus cluster. The phylogenetic tree on the left represents the relationships among genus clusters, indicating with a star the ones for which isolated representatives are available. The labels “*Lg*”, “*Li*”, “*Lo*”, “*Lm*” and “*Lp*” represent the genus clusters including *L. geestiana*, *L. israelensis*, *L. oakridgensis*, *L. micdadei* and *L. pneumophila*, respectively.

### 2.3 *Legionellaceae* species have distinct environmental distributions

Given the genotypic diversity among *Legionellaceae* species, it is expected that they will have distinct, but partly overlapping, ecological niches and environmental distributions. To assess this, we used the OTUs provided by MicrobeAtlas (85), linking them to the full-length 16S rRNA genes of 38 species clusters through reciprocal best hits (Table S11). We detected most *Legionellaceae* OTUs in all major environmental classes (Figure 5), confirming the variety of niches of these bacteria (Figure 1) and highlighting a surprisingly high fraction of animal-associated samples. A closer inspection (Figure S9) revealed several samples from gastrointestinal tract or feces, as well as non-human hosts. On one hand, while the detection of *Legionellaceae* bacteria in feces could result from the presence of these bacteria in the ingested water, previous studies reported of *Legionella*-induced diarrhea, indicating how these bacteria can potentially proliferate in the gut (86). On the other, insects and cattle appear among the non-human hosts category. Insects represent known hosts for intracellular bacteria (i.e., *L. polyplacis*) (87). Cattle, as well as other animals have also been reported as being infected by *Legionellaceae* bacteria (88). Our result highlight how, despite the very limited research regarding the prevalence of *Legionellaceae* bacteria among non-human hosts, such occurrences are likely more common than what previously thought.

**Figure 5.**
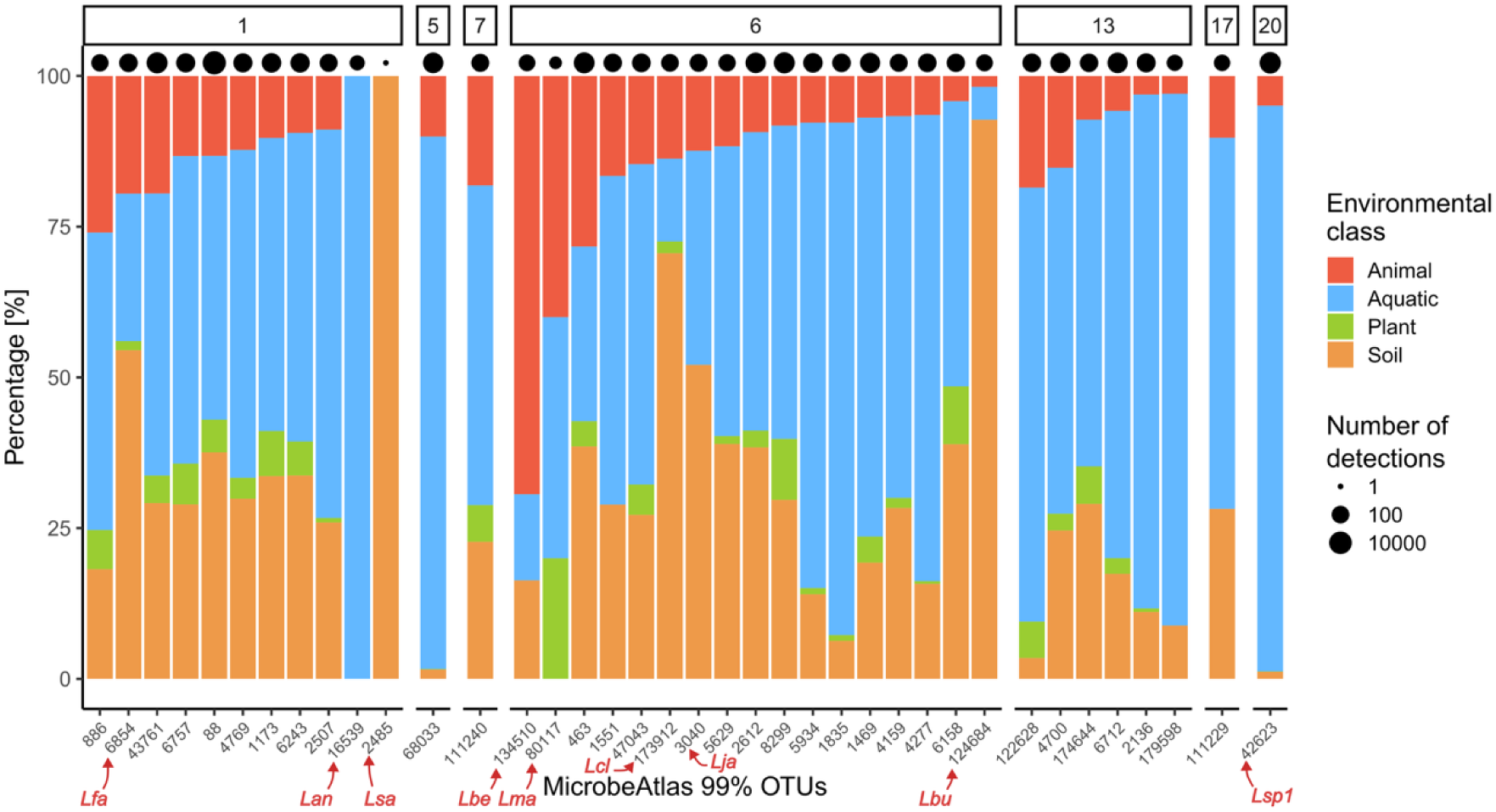
MicrobeAtlas OTUs (99% similarity) matching full-length 16S rRNA from the analyzed genomes, together with the number of samples in which they were found (i.e., OTUs’ occurrence) and their distribution among major environmental classes. Species labels are the following: *L. antarctica (Lan), L. beliardensis (Lbe), L. busanensis (Lbu), L. clemsoniensis (Lcl), L. fallonii (Lfa), L. jamestowniensis (Lja), L. maceachernii (Lma), saoudiensis (Lsa), spp cluster 94_0 (Lsp1)*.

Generally we found OTUs associated to the analyzed genomes mostly in aquatic samples, yet several OTUs showed a drastically higher prevalence of soil samples or animal-associated ones (Figure 5). We did not observe a statistical difference in the occurrence of OTUs belonging to different genus clusters (Kruskal-Wallis test, p-val = 0.76) and we detected only a weak effect of genus cluster membership on the distribution over environmental classes (PERMANOVA, R^2^ = 7.6%, p-val = 0.07), likely caused by the heterogeneity present within these broad environmental classifications. We found more similar environmental distributions for OTUs belonging to the same metabolic cluster (Table S7) than with others (t-test, p-val = 0.04) and at decreasing phylogenetic distance (Spearman correlation, ρ = −0.26, p-val < 0.001). These results highlight that different genus clusters are similarly present in the environment, but their species present distinct, albeit overlapping, ecological niches, corroborating the link between shared evolutionary history, metabolic capabilities and ecological niches.

### 2.4 Implications

Our analysis underscores the extensive genomic diversity within *Legionellaceae*. While key secretion systems (T2SS, T4BSS) are conserved, the family exhibits trait heterogeneity consistent with multiple distinct genera rather than a single genus. The *L. pneumophila* cluster (cluster 1) features larger genomes, lower GC content, and greater metabolic and motility capabilities relative to most other clusters. Similarly, the *L. micdadei* cluster (cluster 6) shows expanded metabolic potential, motility, and T1SS diversity. In contrast, a large separate clade composed primarily of uncultured species (except cluster 13, e.g., *L. oakridgensis*) exhibits reduced genome sizes, restricted motility, and increased auxotrophies. The variability of these traits, as well as variations in clinical relevance and environmental preferences across *Legionellaceae* genus clusters highlight the limitations of maintaining a single genus classification, which conflates major pathogens with low-risk species. Restructuring the family into multiple genera would improve ecological and clinical characterization providing a clearer distinction among the traits in the different genera.. However, given the tight link between taxonomy, detection methods and legislation, such a restructuring should involve all relevant stakeholders is required and requires a joint effort from the research community.

A major knowledge gap remains in the environmental interactions between Legionella and protistan host and how these influence selective pressures over certain traits (e.g., metabolism, secretion systems) and influence environmental prevalence. A better delineation of the environmental niches of clinically-relevant species, as well as yet-to-be-cultured ones will help to identify critical intervention areas to protect public health and provide a more complete picture of the evolution of this family. Such a result, beyond its scientific value, would, allow for better safeguarding public health by allowing the improvement of relevant legislation.

## 3. Materials and Methods

### 3.1 Genomes retrieval, quality assessment and comparison

A total of 5023 genomes classified as belonging to the family *Legionellaceae* were identified from publicly-available sources (date last accession: 15/05/2023, Table S12). Sources included NCBI Assembly database, ENA, JGI and other several large-scale metagenomics survey (89–97). In addition, reads from Wee and coworkers (92) were assembled using spades v3.15.4 (98) after read trimming using fastp v0.23.2 (99) and the removal of known UniVec contaminant (90) using BWA-MEM2 v2.2.1 (100) and samtools v1.16.1 (101). The quality of all the genomes and their taxonomic affiliation was assessed using CheckM2 v1.0.1 (102), and GUNC v1.0.5 (103) and GTDB-Tk v2.3.0 (104) to remove genomes with more than 5% contamination, less than 85% complete and not belonging to the investigated family, eventually after manual refinement using Anvi’o v8 (105). Genomes dereplication was performed with an initial clustering at 90% ANI using MASH v1.1 (106), followed by a secondary one using skani v0.2.1 (107) at 95% and 99.5% ANI. Representative genomes for species (i.e., 95% ANI) and genomovars (i.e., 99.5% ANI) were chosen as the ones with the highest value derived from the following formula (108): *Completeness* – 5\**Contamination* + 0.5*log(*N50*) + *Centrality*/100, using CheckM2 quality estimates and where *N50* is the minimum length for which half of the genome belongs to contigs longer than *N50*, while *Centrality* is the average ANI to genomes within the same secondary cluster. Due to the high number of genomovars found for *L. pneumophila* and *L. fraseri,* only the species representative and the three representatives of the genomovar with the highest number of genomes were selected. Species’ clinical-associations were retrieved from published literature (12).

### 3.2 Proteins predictions and functional annotations

Proteins were predicted using Bakta v1.8.1 (109) (v5.0, full database) and annotated using eggNOG-mapper v2.1.12 (110). Amino acid similarity was assessed using FastAAI v0.1.20 (111), while POCP-nf v2.3.6 was used to calculate the percentage of conserved proteins among genomes (112,113). T4SS were detected using CONJScan v2.0.1 (114), while other secretion systems, flagella and pili using TXSScan v1.1.3 (115). T6SSs were considered as present in case more than nine proteins were detected in order to match previous reports (40). Plasmid-associated contigs were detected using Platon v1.7 (116). Sensory and regulatory proteins were selected based on eggNOG-mapper annotations and the signal transduction specific KEGG profiles from MiST v4.0 database (117). Chemotaxis and methyl-accepting proteins were detected using HMMER v3.4 (118) and the models from the MiST database. KEGG metabolic pathways completeness was estimated using DRAM v1.4.6 (119), while the amino acids biosynthetic potential was evaluated using GapMind and gapseq v1.2. Clusters of genomes presenting similar metabolic capabilities were obtained through HDBSCAN clustering based on the euclidean distance of metabolic pathways completeness. CAZYmes were annotated against the dbCAN3 database using dbcan_run v4.0.0 (120), retaining only proteins identified by at least two detection methods. Peptidases were identified searching the predicted proteins against the MEROPS v12.4 database (121) using MMseqs2 v14-7e284 (122) retaining proteins showing at least 30% identity over 65% query coverage and displaying a bitscore ≥50. Secreted proteins were identified using SignalP v6.0 (123). Homologs of selected *L. pneumophila* strain *Paris* proteins were detected using BlastP v2.12.0 (124) using the same thresholds used for MMseqs2 and inspecting Bakta and eggNOG-mapper annotations (e.g., PFAMs detected). Type 1 secretion systems were considered as present if both LssD/lrpssD/lb1ssD/lb2ssD and LssB/lrpssB/lb1ssB/lb2ssB proteins were detected within a five proteins neighbourhood (50). Eventual duplicated matches with TXSScan results were assessed using a custom script.

### 3.3 Pangenome and phylogeny construction

Predicted proteins were used to perform calculations using PPanGGOLiN v1.2.105 (125) employing a minimum identity percentage of 30% and minimum coverage of 65%, as done previously (54) and achieving high concordance (adjusted rand index = 0.96) with eggNOG orthogroups. Single-copy orthologs were identified using BUSCO v6.0.0 (126) based on the lineage “legionellales_odb12” in all the retained genomes, as well as outgroup genomes originating from other *Legionellales* families (*Coxiella burnetii*: GCF_005280755.1; *C. mudrowiae*: GCF_001077715.1: *Berkiella aquae*: GCF_001431295.2; *B. cookevillensis*: GCF_001431315.2; *Aquicella lusitana*: GCF_902459475.1; *A. siphonis*: GCF_902459485.1). Proteins were aligned individually using MAFFT G-INS-i, and then used to predict a partitioned maximum-likelihood phylogenetic tree using IQ-TREE v2.2.2.6 (127) testing both LG and WAG models using automated model selection, as well as considering mixture model classes and merging partitions with comparable evolutionary rates (parameters: *-mset LG,WAG -m MFP+C+MERGE –cmin 10 – cmax 60*) (128). Support values were calculated based on 1000 ultrafast bootstraps with bootstrap tree optimization (129). The resulting phylogenetic tree was visualized and annotated in iTOL v6 (130).

### 3.4 Assessment of species environmental distribution

One random full length 16S rRNA gene per genome was retrieved per species from the Bakta annotation. Such sequences were matched to the full-length OTUs (99% similarity) retrieved from MicrobeAtlas v3.0 (downloaded on: 16/9/2024) through MMseqs2’s reciprocal best hit workflow and removing OTUs which showed best hits to multiple species clusters. Major environmental classes were obtained from MicrobeAtlas metadata, selecting only samples for which only a single class was listed. Spearman correlations were used to estimate the concordance of the environmental distribution among selected OTUs.

### 3.5 Statistical testing and data analysis

Analysis of the bioinformatic results was performed in R v4.3.0 (131) leveraging the functions provided by libraries Tidyverse v2.0.0 (132), ape v5.7-1 (133), dbscan v1.2-0 (134) and vegan v2.6-4 (135). Statistical significance was assessed using appropriate two-tailed tests and a threshold p-value of 0.05.

## 4. Data availability

Supplementary data and tables S1-S12, as well as the *Legionellaceae* phylogenetic tree are available on Github at the following link: https://github.com/mgabriell1/Legionellaceae_diversity.

## 6. Author contribution statement

MG: Conceptualization, Methodology, Formal Analysis, investigation, Data curation, Writing – Original draft, Visualization; AC: Writing – Review and editing; FH: Writing – Review and Editing, Supervision, Funding acquisition.

## Acknowledgments

This research was funded through the LeCo project (Legionella Control in Buildings; Aramis nr.: 4.20.01) by the Federal Food Safety and Veterinary Office (FSVO), in partnership with the Federal Offices of Public Health (FOPH) and Energy (SFOE) of Switzerland, as well as Eawag discretionary funding. The authors would also like to acknowledge Alyssa Henderson for her constructive feedback on genomic-scale metabolic models, David Johnson and Tim Julian for the valuable comments on an initial manuscript version and Valeria Gaia for providing additional *Legionellaceae* isolates.

## References

1. Thompson LR, Sanders JG, McDonald D, Amir A, Ladau J, Locey KJ, et al. A communal catalogue reveals Earth’s multiscale microbial diversity. Nature. 2017 Nov 23;551(7681):457–63. doi:10.1038/nature24621

2. Blount ZD. The unexhausted potential of E. coli. eLife. 2015 Mar 25;4:e05826. doi:10.7554/eLife.05826

3. Hrovat K, Dutilh BE, Medema MH, Melkonian C. Taxonomic resolution of different 16S rRNA variable regions varies strongly across plant-associated bacteria. ISME Communications. 2024 Jan 8;4(1):ycae034. doi:10.1093/ismeco/ycae034

4. Leigh JA, Albers SV, Atomi H, Allers T. Model organisms for genetics in the domain Archaea: methanogens, halophiles, Thermococcales and Sulfolobales. FEMS Microbiol Rev. 2011 Jul;35(4):577–608. doi:10.1111/j.1574-6976.2011.00265.x

5. Beukes CW, Palmer M, Manyaka P, Chan WY, Avontuur JR, Van Zyl E, et al. Genome Data Provides High Support for Generic Boundaries in Burkholderia Sensu Lato. Front Microbiol. 2017 Jun 26;8:1154. doi:10.3389/fmicb.2017.01154

6. Van Rossum T, Ferretti P, Maistrenko OM, Bork P. Diversity within species: interpreting strains in microbiomes. Nat Rev Microbiol. 2020 Sep;18(9):491–506. doi:10.1038/s41579-020-0368-1

7. Crosby KC, Rojas M, Sharma P, Johnson MA, Mazloom R, Kvitko BH, et al. Genomic delineation and description of species and within-species lineages in the genus Pantoea. Front Microbiol. 2023 Nov 9;14:1254999. doi:10.3389/fmicb.2023.1254999

8. Baker-Goering M, Roy K, Edens C, Collier S. Economic Burden of Legionnaires’ Disease, United States, 2014. Emerg Infect Dis. 2021 Jan;27(1):255–7. doi:10.3201/eid2701.191198

9. Moffa MA, Rock C, Galiatsatos P, Gamage SD, Schwab KJ, Exum NG. Legionellosis on the rise: A scoping review of sporadic, community-acquired incidence in the United States. Epidemiol Infect. 2023;151:e133. doi:10.1017/S0950268823001206

10. LPSN. Family Legionellaceae [Internet]. [cited 2024 Dec 8]. Available from: https://lpsn.dsmz.de/family/legionellaceae

11. Yu VL, Plouffe JF, Pastoris MC, Stout JE, Schousboe M, Widmer A, et al. Distribution of Legionella Species and Serogroups Isolated by Culture in Patients with Sporadic Community-Acquired Legionellosis: An International Collaborative Survey. J INFECT DIS. 2002 Jul;186(1):127–8. doi:10.1086/341087

12. LeCo Project. LegioSpecies - an updated and referenced list of Legionellaceae species [Internet]. 2024 [cited 2024 Oct 11]. Available from: https://zenodo.org/doi/10.5281/zenodo.11072744doi:10.5281/ZENODO.11072744

13. van der Wielen PWJJ, Wierenga W, Oesterholt FIHM, Oostdijk A, van der Werff A. Rightly to an effective Legionella prevention (in Dutch). Utrecht, The Netherlands; 2021.

14. National Academies of Sciences, Engineering, and Medicine. Management of Legionella in Water Systems [Internet]. Washington, D.C.: National Academies Press; 2020 [cited 2024 Oct 11]. Available from: https://www.nap.edu/catalog/25474doi:10.17226/25474

15. Van Kenhove E, Dinne K, Janssens A, Laverge J. Overview and comparison of Legionella regulations worldwide. American Journal of Infection Control. 2019 Aug;47(8):968–78. doi:10.1016/j.ajic.2018.10.006

16. Joseph SJ, Cox D, Wolff B, Morrison SS, Kozak-Muiznieks NA, Frace M, et al. Dynamics of genome change among Legionella species. Sci Rep. 2016 Sep 16;6(1):33442. doi:10.1038/srep33442

17. Gomez-Valero L, Rusniok C, Carson D, Mondino S, Pérez-Cobas AE, Rolando M, et al. More than 18,000 effectors in the Legionella genus genome provide multiple, independent combinations for replication in human cells. Proc Natl Acad Sci USA. 2019 Feb 5;116(6):2265–73. doi:10.1073/pnas.1808016116

18. Tobias NJ, Ahrendt T, Schell U, Miltenberger M, Hilbi H, Bode HB. Legionella shows a diverse secondary metabolism dependent on a broad spectrum Sfp-type phosphopantetheinyl transferase. PeerJ. 2016 Nov 24;4:e2720. doi:10.7717/peerj.2720

19. Gomez-Valero L, Chiner-Oms A, Comas I, Buchrieser C. Evolutionary Dissection of the Dot/Icm System Based on Comparative Genomics of 58 Legionella Species. Hershberg R, editor. Genome Biology and Evolution. 2019 Sep 1;11(9):2619–32. doi:10.1093/gbe/evz186

20. White RC, Cianciotto NP. Assessing the impact, genomics and evolution of type II secretion across a large, medically important genus: the Legionella type II secretion paradigm. Microbial Genomics. 2019 Jun 1;5(6). doi:10.1099/mgen.0.000273

21. Graells T, Ishak H, Larsson M, Guy L. The all-intracellular order Legionellales is unexpectedly diverse, globally distributed and lowly abundant. FEMS Microbiology Ecology. 2018 Dec 1;94(12). doi:10.1093/femsec/fiy185

22. Vartoukian SR, Palmer RM, Wade WG. Strategies for culture of ‘unculturable’ bacteria: Culturing the unculturable. FEMS Microbiology Letters. 2010;309:1–7. doi:10.1111/j.1574-6968.2010.02000.x

23. Rodriguez-R LM, Conrad RE, Viver T, Feistel DJ, Lindner BG, Venter SN, et al. An ANI gap within bacterial species that advances the definitions of intra-species units. Jouline IB, editor. mBio. 2024 Jan 16;15(1):e02696–23. doi:10.1128/mbio.02696-23

24. Oren A, Garrity G. Notification of changes in taxonomic opinion previously published outside the IJSEM. List of changes in taxonomic opinion no. 35. International Journal of Systematic and Evolutionary Microbiology. 2022 Feb 3;72(1). doi:10.1099/ijsem.0.005164

25. Saini N, Gupta RS. A robust phylogenetic framework for members of the order Legionellales and its main genera (Legionella, Aquicella, Coxiella and Rickettsiella) based on phylogenomic analyses and identification of molecular markers demarcating different clades. Antonie van Leeuwenhoek. 2021 Jul;114(7):957–82. doi:10.1007/s10482-021-01569-9

26. Kozak-Muiznieks NA, Morrison SS, Mercante JW, Ishaq MK, Johnson T, Caravas J, et al. Comparative genome analysis reveals a complex population structure of Legionella pneumophila subspecies. Infection, Genetics and Evolution. 2018 Apr;59:172–85. doi:10.1016/j.meegid.2018.02.008

27. Philippot L, Andersson SGE, Battin TJ, Prosser JI, Schimel JP, Whitman WB, et al. The ecological coherence of high bacterial taxonomic ranks. Nat Rev Microbiol. 2010 Jul;8(7):523–9. doi:10.1038/nrmicro2367

28. Konstantinidis KT, Tiedje JM. Prokaryotic taxonomy and phylogeny in the genomic era: advancements and challenges ahead. Current Opinion in Microbiology. 2007 Oct;10(5):504–9. doi:10.1016/j.mib.2007.08.006

29. Qin QL, Xie BB, Zhang XY, Chen XL, Zhou BC, Zhou J, et al. A Proposed Genus Boundary for the Prokaryotes Based on Genomic Insights. J Bacteriol. 2014 Jun 15;196(12):2210–5. doi:10.1128/JB.01688-14

30. Chambers ST, Slow S, Scott-Thomas A, Murdoch DR. Legionellosis Caused by Non-Legionella pneumophila Species, with a Focus on Legionella longbeachae. Microorganisms. 2021 Jan 31;9(2):291. doi:10.3390/microorganisms9020291

31. Muder RR, Yu VL. Infection Due to Legionella Species Other Than L. pneumophila. Clinical Infectious Diseases. 2002 Oct 15;35(8):990–8. doi:10.1086/342884

32. Wang C, Chuai X, Liang M. Legionella feeleii: pneumonia or Pontiac fever? Bacterial virulence traits and host immune response. Med Microbiol Immunol. 2019 Feb;208(1):25–32. doi:10.1007/s00430-018-0571-0

33. Mahajan S, Agashe D. Evolutionary jumps in bacterial GC content. Wong A, editor. G3 Genes|Genomes|Genetics. 2022 Jul 29;12(8):jkac108. doi:10.1093/g3journal/jkac108

34. Santos-Garcia D, Morel O, Henri H, El Filali A, Buysse M, Noël V, et al. Genomic changes during the evolution of the Coxiella genus along the parasitism-mutualism continuum. Peer Community Journal. 2023 May 2;3:e41. doi:10.24072/pcjournal.269

35. Köstlbacher S, Collingro A, Halter T, Schulz F, Jungbluth SP, Horn M. Pangenomics reveals alternative environmental lifestyles among chlamydiae. Nat Commun. 2021 Jun 29;12(1):4021. doi:10.1038/s41467-021-24294-3

36. Tettelin H, Riley D, Cattuto C, Medini D. Comparative genomics: the bacterial pan-genome. Current Opinion in Microbiology. 2008 Oct;11(5):472–7. doi:10.1016/j.mib.2008.09.006

37. Poirier V, Av-Gay Y. Intracellular Growth of Bacterial Pathogens: The Role of Secreted Effector Proteins in the Control of Phagocytosed Microorganisms. Kudva IT, Bannantine JP, editors. Microbiol Spectr. 2015 Dec 18;3(6):3.6.12. doi:10.1128/microbiolspec.VMBF-0003-2014

38. Wallden K, Rivera-Calzada A, Waksman G. Microreview: Type IV secretion systems: versatility and diversity in function: Diversity and versatility of the type IV secretion system. Cellular Microbiology. 2010 Sep;12(9):1203–12. doi:10.1111/j.1462-5822.2010.01499.x

39. Øíhová J, Nováková E, Husník F, Hypša V. Legionella Becoming a Mutualist: Adaptive Processes Shaping the Genome of Symbiont in the Louse Polyplax serrata. Genome Biology and Evolution. 2017 Nov 1;9(11):2946–57. doi:10.1093/gbe/evx217

40. Qin T, Zhou H, Ren H, Liu W. Distribution of Secretion Systems in the Genus Legionella and Its Correlation with Pathogenicity. Front Microbiol. 2017 Mar 14;08. doi:10.3389/fmicb.2017.00388

41. Söderberg MA, Dao J, Starkenburg SR, Cianciotto NP. Importance of Type II Secretion for Survival of Legionella pneumophila in Tap Water and in Amoebae at Low Temperatures. Appl Environ Microbiol. 2008 Sep;74(17):5583–8. doi:10.1128/AEM.00067-08

42. Carrión VJ, Perez-Jaramillo J, Cordovez V, Tracanna V, De Hollander M, Ruiz-Buck D, et al. Pathogen-induced activation of disease-suppressive functions in the endophytic root microbiome. Science. 2019 Nov;366(6465):606–12. doi:10.1126/science.aaw9285

43. Wang L, Liu W, Liang J, Zhao L, Li Q, Zhou C, et al. Mining of novel secondary metabolite biosynthetic gene clusters from acid mine drainage. Sci Data. 2022 Dec 9;9(1):760. doi:10.1038/s41597-022-01866-6

44. De Buck E, Anné J, Lammertyn E. The role of protein secretion systems in the virulence of the intracellular pathogen Legionella pneumophila. Microbiology. 2007 Dec 1;153(12):3948–53. doi:10.1099/mic.0.2007/012039-0

45. Gomez-Valero L, Rusniok C, Rolando M, Neou M, Dervins-Ravault D, Demirtas J, et al. Comparative analyses of Legionella species identifies genetic features of strains causing Legionnaires!] disease. Genome Biology. 2014;15(505). doi:10.1186/s13059-014-0505-0

46. Fuche F, Vianney A, Andrea C, Doublet P, Gilbert C. Functional Type 1 Secretion System Involved in Legionella pneumophila Virulence. Parkinson JS, editor. J Bacteriol. 2015 Feb 1;197(3):563–71. doi:10.1128/JB.02164-14

47. Meuskens I, Saragliadis A, Leo JC, Linke D. Type V Secretion Systems: An Overview of Passenger Domain Functions. Front Microbiol. 2019 May 31;10:1163. 2019 May 31;10:1163. doi:. doi:10.3389/fmicb.2019.01163

48. Gallegos-Monterrosa R, Coulthurst SJ. The ecological impact of a bacterial weapon: microbial interactions and the Type VI secretion system. FEMS Microbiology Reviews. 2021 Nov 23;45(6):fuab033. doi:10.1093/femsre/fuab033

49. Stubbusch AKM, Peaudecerf FJ, Lee KS, Paoli L, Schwartzman J, Stocker R, et al. Antagonism as a foraging strategy in microbial communities. Science. 2025 Jun 12;388(6752):1214–7. doi:10.1126/science.adr8286

50. Brown CL, Garner E, Jospin G, Coil DA, Schwake DO, Eisen JA, et al. Whole genome sequence analysis reveals the broad distribution of the RtxA type 1 secretion system and four novel putative type 1 secretion systems throughout the Legionella genus. Voth DE, editor. PLoS ONE. 2020 Jan 14;15(1):e0223033. doi:10.1371/journal.pone.0223033

51. Brown AW, Simone PS, York JC, Emmert GL. A device for fully automated on-site process monitoring and control of trihalomethane concentrations in drinking water. Analytica Chimica Acta. 2015 Jan;853:351–9. doi:10.1016/j.aca.2014.10.052

52. Qin T, Zhou H, Ren H, Liu W. Distribution of Secretion Systems in the Genus Legionella and Its Correlation with Pathogenicity. Front Microbiol. 2017 Mar 14;08. doi:10.3389/fmicb.2017.00388

53. Ensminger AW. Legionella pneumophila, armed to the hilt: justifying the largest arsenal of effectors in the bacterial world. Current Opinion in Microbiology. 2016 Feb;29:74–80. doi:10.1016/j.mib.2015.11.002

54. Gomez-Valero L, Rusniok C, Carson D, Mondino S, Pérez-Cobas AE, Rolando M, et al. More than 18,000 effectors in the Legionella genus genome provide multiple, independent combinations for replication in human cells. Proc Natl Acad Sci USA. 2019 Feb 5;116(6):2265–73. doi:10.1073/pnas.1808016116

55. Raina JB, Fernandez V, Lambert B, Stocker R, Seymour JR. The role of microbial motility and chemotaxis in symbiosis. Nat Rev Microbiol. 2019 May;17(5):284–94. doi:10.1038/s41579-019-0182-9

56. Shaheen M, Ashbolt NJ. Differential Bacterial Predation by Free-Living Amoebae May Result in Blooms of Legionella in Drinking Water Systems. Microorganisms. 2021 Jan 15;9(1):174. doi:10.3390/microorganisms9010174

57. Duncan C, Prashar A, So J, Tang P, Low DE, Terebiznik M, et al. Lcl of Legionella pneumophila Is an Immunogenic GAG Binding Adhesin That Promotes Interactions with Lung Epithelial Cells and Plays a Crucial Role in Biofilm Formation. Weiser JN, editor. Infect Immun. 2011 Jun;79(6):2168–81. doi:10.1128/IAI.01304-10

58. Matilla MA, Krell T. The effect of bacterial chemotaxis on host infection and pathogenicity. FEMS Microbiology Reviews. 2018 Jan 1;42(1). doi:10.1093/femsre/fux052

59. Dietrich C, Heuner K, Brand BC, Hacker J, Steinert M. Flagellum of Legionella pneumophila Positively Affects the Early Phase of Infection of Eukaryotic Host Cells. DiRita VJ, editor. Infect Immun. 2001 Apr;69(4):2116–22. doi:10.1128/IAI.69.4.2116-2122.2001

60. Wuichet K, Zhulin IB. Origins and Diversification of a Complex Signal Transduction System in Prokaryotes. Sci Signal. 2010 Jun 29;3(128). doi:10.1126/scisignal.2000724

61. Seymour JR, Brumley DR, Stocker R, Raina JB. Swimming towards each other: the role of chemotaxis in bacterial interactions. Trends in Microbiology. 2024 Jul;32(7):640–9. doi:10.1016/j.tim.2023.12.008

62. Coil DA, Anné J. Twitching motility in Legionella pneumophila. FEMS Microbiology Letters. 2009 Apr;293(2):271–7. doi:10.1111/j.1574-6968.2009.01532.x

63. Oliva G, Sahr T, Buchrieser C. The Life Cycle of L. pneumophila: Cellular Differentiation Is Linked to Virulence and Metabolism. Front Cell Infect Microbiol. 2018 Jan 19;8:3. doi:10.3389/fcimb.2018.00003

64. Eylert E, Herrmann V, Jules M, Gillmaier N, Lautner M, Buchrieser C, et al. Isotopologue Profiling of Legionella pneumophila. Journal of Biological Chemistry. 2010 Jul;285(29):22232–43. doi:10.1074/jbc.M110.128678

65. Eisenreich W, Heuner K. The life stage-specific pathometabolism of Legionella pneumophila. FEBS Letters. 2016 Nov;590(21):3868–86. doi:10.1002/1873-3468.12326

66. Häuslein I, Manske C, Goebel W, Eisenreich W, Hilbi H. Pathway analysis using ^13^ C-glycerol and other carbon tracers reveals a bipartite metabolism of Legionella pneumophila. Molecular Microbiology. 2016 Apr;100(2):229–46. doi:10.1111/mmi.13313

67. James BW, Mauchline WS, Dennis PJ, Keevil CW, Wait R. Poly-3-Hydroxybutyrate in Legionella pneumophila, an Energy Source for Survival in Low-Nutrient Environments. Appl Environ Microbiol. 1999 Feb;65(2):822–7. doi:10.1128/AEM.65.2.822-827.1999

68. Manske C, Schell U, Hilbi H. Metabolism of myo -Inositol by Legionella pneumophila Promotes Infection of Amoebae and Macrophages. Spormann AM, editor. Appl Environ Microbiol. 2016 Aug 15;82(16):5000–14. doi:10.1128/AEM.01018-16

69. Eisenhofer R, Odriozola I, Alberdi A. Impact of microbial genome completeness on metagenomic functional inference. ISME Communications. 2023 Dec 1;3(1):12. doi:10.1038/s43705-023-00221-z

70. Wilkins AA, Schwarz B, Torres-Escobar A, Castore R, Landry L, Latimer B, et al. The intracellular growth of the vacuolar pathogen Legionella pneumophila is dependent on the acyl chain composition of host membranes. Front Bacteriol. 2024 Feb 12;3:1322138. doi:10.3389/fbrio.2024.1322138

71. Conover GM, Martinez-Morales F, Heidtman MI, Luo ZQ, Tang M, Chen C, et al. Phosphatidylcholine synthesis is required for optimal function of Legionella pneumophila virulence determinants. Cell Microbiol. 2007 Nov 2;0(0):071103031556001-??? doi:10.1111/j.1462-5822.2007.01066.x

72. Eydallin G, Montero M, Almagro G, Sesma MT, Viale AM, Munoz FJ, et al. Genome-Wide Screening of Genes Whose Enhanced Expression Affects Glycogen Accumulation in Escherichia coli. DNA Research. 2010 Apr 1;17(2):61–71. doi:10.1093/dnares/dsp028

73. Lopez-Vazquez CM, Oehmen A, Hooijmans CM, Brdjanovic D, Gijzen HJ, Yuan Z, et al. Modeling the PAO–GAO competition: Effects of carbon source, pH and temperature. Water Research. 2009 Feb;43(2):2. doi:10.1016/j.watres.2008.10.032

74. Nguyen TTH, Myrold DD, Mueller RS. Distributions of Extracellular Peptidases Across Prokaryotic Genomes Reflect Phylogeny and Habitat. Front Microbiol. 2019 Mar 5;10:413. doi:10.3389/fmicb.2019.00413

75. Kadouri D, Jurkevitch E, Okon Y, Castro-Sowinski S. Ecological and Agricultural Significance of Bacterial Polyhydroxyalkanoates. Critical Reviews in Microbiology. 2005 Jan;31(2):55–67. doi:10.1080/10408410590899228

76. McKinlay JB. Are Bacteria Leaky? Mechanisms of Metabolite Externalization in Bacterial Cross-Feeding. Annu Rev Microbiol. 2023 Sep 15;77(1):277–97. doi:10.1146/annurev-micro-032521-023815

77. Ramoneda J, Jensen TBN, Price MN, Casamayor EO, Fierer N. Taxonomic and environmental distribution of bacterial amino acid auxotrophies. Nat Commun. 2023 Nov 22;14(1):7608. doi:10.1038/s41467-023-43435-4

78. D’Souza G, Waschina S, Pande S, Bohl K, Kaleta C, Kost C. Less is more: selective advantages can explain the prevalent loss of biosynthetic genes in bacteria: adaptive loss of biosynthetic genes in bacteria. Evolution. 2014 Sep;68(9):2559–70. doi:10.1111/evo.12468

79. Price MN, Deutschbauer AM, Arkin AP. GapMind: Automated Annotation of Amino Acid Biosynthesis. Hallam SJ, editor. mSystems. 2020 Jun 30;5(3):10.1128/msystems.00291-20. doi:10.1128/msystems.00291-20

80. Zimmermann J, Kaleta C, Waschina S. gapseq: informed prediction of bacterial metabolic pathways and reconstruction of accurate metabolic models. Genome Biol. 2021 Dec;22(1):81. doi:10.1186/s13059-021-02295-1

81. Stipanuk MH. SULFUR AMINO ACID METABOLISM: Pathways for Production and Removal of Homocysteine and Cysteine. Annu Rev Nutr. 2004 Jul 14;24(1):539–77. doi:10.1146/annurev.nutr.24.012003.132418

82. Kredich NM. Biosynthesis of Cysteine. Stewart V, editor. EcoSal Plus. 2008 Jan 12;3(1):10.1128/ecosalplus.3.6.1.11. doi:10.1128/ecosalplus.3.6.1.11

83. George JR, Pine L, Reeves MW, Harrell WK. Amino acid requirements of Legionella pneumophila. J Clin Microbiol. 1980 Mar;11(3):286–91. doi:10.1128/jcm.11.3.286-291.1980

84. Tesh MJ, Miller RD. Amino acid requirements for Legionella pneumophila growth. J Clin Microbiol. 1981 May;13(5):865–9. doi:10.1128/jcm.13.5.865-869.1981

85. Matias Rodrigues JF, Schmidt TSB, Tackmann J, Von Mering C. MAPseq: highly efficient k-mer search with confidence estimates, for rRNA sequence analysis. Birol I, editor. Bioinformatics. 2017 Dec 1;33(23):3808–10. doi:10.1093/bioinformatics/btx517

86. Emuron D, Senadhi V. Gastrointestinal Legionella: An Underecognized Entity and Cause of Secretory Diarrhea: A Case and Review of the Literature: 889. Official journal of the American College of Gastroenterology | ACG [Internet]. 2011;106. Available from: https://journals.lww.com/ajg/fulltext/2011/10002/gastrointestinal_legionella__an_underecognized.889.aspx

87. Kikuchi Y. Endosymbiotic Bacteria in Insects: Their Diversity and Culturability. Microb Environ. 2009;24(3):195–204. doi:10.1264/jsme2.ME09140S

88. Collins MT, Cho SN, Reif JS. Prevalence of antibodies to Legionella pneumophila in animal populations. J Clin Microbiol. 1982 Jan;15(1):130–6. doi:10.1128/jcm.15.1.130-136.1982

89. Chen IMA, Chu K, Palaniappan K, Pillay M, Ratner A, Huang J, et al. IMG/M v.5.0: an integrated data management and comparative analysis system for microbial genomes and microbiomes. Nucleic Acids Research. 2019 Jan 8;47(D1):D666–77. doi:10.1093/nar/gky901

90. Clark K, Karsch-Mizrachi I, Lipman DJ, Ostell J, Sayers EW. GenBank. Nucleic Acids Res. 2016 Jan 4;44(D1):D1. doi:10.1093/nar/gkv1276

91. Leinonen R, Sugawara H, Shumway M, on behalf of the International Nucleotide Sequence Database Collaboration. The Sequence Read Archive. Nucleic Acids Research. 2011 Jan 1;39(Database):Database. doi:10.1093/nar/gkq1019

92. Wee BA, Alves J, Lindsay DSJ, Klatt AB, Sargison FA, Cameron RL, et al. Population analysis of Legionella pneumophila reveals a basis for resistance to complement-mediated killing. Nat Commun. 2021 Dec 9;12(1):7165. doi:10.1038/s41467-021-27478-z

93. Buck M, Garcia SL, Fernandez L, Martin G, Martinez-Rodriguez GA, Saarenheimo J, et al. Comprehensive dataset of shotgun metagenomes from oxygen stratified freshwater lakes and ponds. Sci Data. 2021 May 14;8(1):131. doi:10.1038/s41597-021-00910-1

94. Gurbich TA, Almeida A, Beracochea M, Burdett T, Burgin J, Cochrane G, et al. MGnify Genomes: A Resource for Biome-specific Microbial Genome Catalogues. Journal of Molecular Biology. 2023 Feb;168016. doi:10.1016/j.jmb.2023.168016

95. Nayfach S, Roux S, Seshadri R, Udwary D, Varghese N, Schulz F, et al. A genomic catalog of Earth’s microbiomes. Nat Biotechnol. 2021 Apr;39(4):499–509. doi:10.1038/s41587-020-0718-6

96. Nishimura Y, Yoshizawa S. The OceanDNA MAG catalog contains over 50,000 prokaryotic genomes originated from various marine environments. Sci Data. 2022 Jun 17;9(1):305. doi:10.1038/s41597-022-01392-5

97. Sudarshan AS, Dai Z, Gabrielli M, Oosthuizen-Vosloo S, Konstantinidis KT, Pinto AJ. New Drinking Water Genome Catalog Identifies a Globally Distributed Bacterial Genus Adapted to Disinfected Drinking Water Systems. Environ Sci Technol. 2024 Sep 17;58(37):16475–87. doi:10.1021/acs.est.4c05086

98. Prjibelski A, Antipov D, Meleshko D, Lapidus A, Korobeynikov A. Using SPAdes De Novo Assembler. CP in Bioinformatics. 2020 Jun;70(1):e102. doi:10.1002/cpbi.102

99. Chen S, Zhou Y, Chen Y, Gu J. fastp: an ultra-fast all-in-one FASTQ preprocessor. Bioinformatics. 2018 Sep 1;34(17):i884–90. doi:10.1093/bioinformatics/bty560

100. Vasimuddin Md, Misra S, Li H, Aluru S. Efficient Architecture-Aware Acceleration of BWA-MEM for Multicore Systems. In: 2019 IEEE International Parallel and Distributed Processing Symposium (IPDPS) [Internet]. Rio de Janeiro, Brazil: IEEE; 2019 [cited 2022 Oct 11]. p. 314–24. Available from: https://ieeexplore.ieee.org/document/8820962/ doi:10.1109/IPDPS.2019.00041

101. Li H, Handsaker B, Wysoker A, Fennell T, Ruan J, Homer N, et al. The Sequence Alignment/Map format and SAMtools. Bioinformatics. 2009 Aug 15;25(16):16. doi:10.1093/bioinformatics/btp352

102. Chklovski A, Parks DH, Woodcroft BJ, Tyson GW. CheckM2: a rapid, scalable and accurate tool for assessing microbial genome quality using machine learning. Nat Methods. 2023 Aug;20(8):1203–12. doi:10.1038/s41592-023-01940-w

103. Orakov A, Fullam A, Coelho LP, Khedkar S, Szklarczyk D, Mende DR, et al. GUNC: detection of chimerism and contamination in prokaryotic genomes. Genome Biol. 2021 Dec;22(1):178. doi:10.1186/s13059-021-02393-0

104. Chaumeil PA, Mussig AJ, Hugenholtz P, Parks DH. GTDB-Tk v2: memory friendly classification with the genome taxonomy database. Borgwardt K, editor. Bioinformatics. 2022 Nov 30;38(23):5315–6. doi:10.1093/bioinformatics/btac672

105. Eren AM, Kiefl E, Shaiber A, Veseli I, Miller SE, Schechter MS, et al. Community-led, integrated, reproducible multi-omics with anvi’o. Nat Microbiol. 2020 Dec 21;6(1):3–6. doi:10.1038/s41564-020-00834-3

106. Ondov BD, Treangen TJ, Melsted P, Mallonee AB, Bergman NH, Koren S, et al. Mash: fast genome and metagenome distance estimation using MinHash. Genome Biol. 2016 Dec;17(1):1. doi:10.1186/s13059-016-0997-x

107. Shaw J, Yu YW. Fast and robust metagenomic sequence comparison through sparse chaining with skani. Nat Methods. 2023 Nov;20(11):1661–5. doi:10.1038/s41592-023-02018-3

108. Olm MR, Brown CT, Brooks B, Banfield JF. dRep: a tool for fast and accurate genomic comparisons that enables improved genome recovery from metagenomes through de-replication. ISME J. 2017 Dec;11(12):2864–8. doi:10.1038/ismej.2017.126

109. Schwengers O, Jelonek L, Dieckmann MA, Beyvers S, Blom J, Goesmann A. Bakta: rapid and standardized annotation of bacterial genomes via alignment-free sequence identification: Find out more about Bakta, the motivation, challenges and applications, here. Microbial Genomics. 2021 Nov 30;7(11). doi:10.1099/mgen.0.000685

110. Cantalapiedra CP, Hernández-Plaza A, Letunic I, Bork P, Huerta-Cepas J. eggNOG-mapper v2: Functional Annotation, Orthology Assignments, and Domain Prediction at the Metagenomic Scale. Tamura K, editor. Molecular Biology and Evolution. 2021 Dec 9;38(12):5825–9. doi:10.1093/molbev/msab293

111. Konstantinidis K, Ruiz-Perez C, Gerhardt K, Rodriguez-R L, Jain C, Tiedje J, et al. FastAAI: Efficient Estimation of Genome Average Amino Acid Identity and Phylum-level relationships using Tetramers of Universal Proteins [Internet]. 2022 [cited 2024 Oct 11]. Available from: https://www.researchsquare.com/article/rs-1459378/v1 doi:10.21203/rs.3.rs-1459378/v1

112. Hölzer M. POCP-nf: an automatic Nextflow pipeline for calculating the percentage of conserved proteins in bacterial taxonomy. Martelli PL, editor. Bioinformatics. 2024 Mar 29;40(4):btae175. doi:10.1093/bioinformatics/btae175

113. Buchfink B, Reuter K, Drost HG. Sensitive protein alignments at tree-of-life scale using DIAMOND. Nat Methods. 2021 Apr;18(4):4. doi:10.1038/s41592-021-01101-x

114. Cury J, Abby SS, Doppelt-Azeroual O, Néron B, Rocha EPC. Identifying Conjugative Plasmids and Integrative Conjugative Elements with CONJscan. In: De La Cruz F, editor. Horizontal Gene Transfer [Internet]. New York, NY: Springer US; 2020 [cited 2024 Oct 11]. p. 265–83. (Methods in Molecular Biology). Available from: http://link.springer.com/10.1007/978-1-4939-9877-7_19 doi:10.1007/978-1-4939-9877-7_19

115. Abby SS, Cury J, Guglielmini J, Néron B, Touchon M, Rocha EPC. Identification of protein secretion systems in bacterial genomes. Sci Rep. 2016 Mar 16;6(1):23080. doi:10.1038/srep23080

116. Schwengers O, Barth P, Falgenhauer L, Hain T, Chakraborty T, Goesmann A. Platon: identification and characterization of bacterial plasmid contigs in short-read draft assemblies exploiting protein sequence-based replicon distribution scores. Microbial Genomics. 2020 Oct 1;6(10). doi:10.1099/mgen.0.000398

117. Gumerov VM, Ulrich LE, Zhulin IB. MiST 4.0: a new release of the microbial signal transduction database, now with a metagenomic component. Nucleic Acids Research. 2024 Jan 5;52(D1):D647–53. doi:10.1093/nar/gkad847

118. Eddy SR. Accelerated Profile HMM Searches. Pearson WR, editor. PLoS Comput Biol. 2011 Oct 20;7(10):e1002195. doi:10.1371/journal.pcbi.1002195

119. Shaffer M, Borton MA, McGivern BB, Zayed AA, La Rosa SL, Solden LM, et al. DRAM for distilling microbial metabolism to automate the curation of microbiome function. Nucleic Acids Research. 2020 Sep 18;48(16):8883–900. doi:10.1093/nar/gkaa621

120. Zheng J, Ge Q, Yan Y, Zhang X, Huang L, Yin Y. dbCAN3: automated carbohydrate-active enzyme and substrate annotation. Nucleic Acids Research. 2023 Jul 5;51(W1):W115–21. doi:10.1093/nar/gkad328

121. Rawlings ND, Barrett AJ, Thomas PD, Huang X, Bateman A, Finn RD. The MEROPS database of proteolytic enzymes, their substrates and inhibitors in 2017 and a comparison with peptidases in the PANTHER database. Nucleic Acids Research. 2018 Jan 4;46(D1):D624–32. doi:10.1093/nar/gkx1134

122. Steinegger M, Söding J. MMseqs2 enables sensitive protein sequence searching for the analysis of massive data sets. Nat Biotechnol. 2017 Nov;35(11):1026–8. doi:10.1038/nbt.3988

123. Teufel F, Almagro Armenteros JJ, Johansen AR, Gíslason MH, Pihl SI, Tsirigos KD, et al. SignalP 6.0 predicts all five types of signal peptides using protein language models. Nat Biotechnol. 2022 Jul;40(7):1023–5. doi:10.1038/s41587-021-01156-3

124. Camacho C, Coulouris G, Avagyan V, Ma N, Papadopoulos J, Bealer K, et al. BLAST+: architecture and applications. BMC Bioinformatics. 2009 Dec;10(1):421. doi:10.1186/1471-2105-10-421

125. Gautreau G, Bazin A, Gachet M, Planel R, Burlot L, Dubois M, et al. PPanGGOLiN: Depicting microbial diversity via a partitioned pangenome graph. Ouzounis CA, editor. PLoS Comput Biol. 2020 Mar 19;16(3):e1007732. doi:10.1371/journal.pcbi.1007732

126. Tegenfeldt F, Kuznetsov D, Manni M, Berkeley M, Zdobnov EM, Kriventseva EV. OrthoDB and BUSCO update: annotation of orthologs with wider sampling of genomes. Nucleic Acids Research. 2025 Jan 6;53(D1):D516–22. doi:10.1093/nar/gkae987

127. Minh BQ, Schmidt HA, Chernomor O, Schrempf D, Woodhams MD, Von Haeseler A, et al. IQ-TREE 2: New Models and Efficient Methods for Phylogenetic Inference in the Genomic Era. Teeling E, editor. Molecular Biology and Evolution. 2020 May 1;37(5):1530–4. doi:10.1093/molbev/msaa015

128. Kalyaanamoorthy S, Minh BQ, Wong TKF, Von Haeseler A, Jermiin LS. ModelFinder: fast model selection for accurate phylogenetic estimates. Nat Methods. 2017 Jun;14(6):587–9. doi:10.1038/nmeth.4285

129. Hoang DT, Chernomor O, Von Haeseler A, Minh BQ, Vinh LS. UFBoot2: Improving the Ultrafast Bootstrap Approximation. Molecular Biology and Evolution. 2018 Feb 1;35(2):518–22. doi:10.1093/molbev/msx281

130. Letunic I, Bork P. Interactive Tree of Life (iTOL) v6: recent updates to the phylogenetic tree display and annotation tool. Nucleic Acids Research. 2024 Jul 5;52(W1):W78–82. doi:10.1093/nar/gkae268

131. R Core Team. R: A Language and Environment for Statistical Computing [Internet]. Vienna, Austria: R Foundation for Statistical Computing; 2023. Available from: https://www.R-project.org/

132. Wickham H, Averick M, Bryan J, Chang W, McGowan L, François R, et al. Welcome to the Tidyverse. JOSS. 2019 Nov 21;4(43):1686. doi:10.21105/joss.01686

133. Paradis E, Claude J, Strimmer K. APE: Analyses of Phylogenetics and Evolution in R language. Bioinformatics. 2004 Jan 22;20(2):289–90. doi:10.1093/bioinformatics/btg412

134. Hahsler M, Piekenbrock M, Doran D. dbscan!]: Fast Density-Based Clustering with R. J Stat Soft. 2019;91(1). doi:10.18637/jss.v091.i01

135. Oksanen J, Simpson GL, Blanchet FG, Kindt R, Legendre P, Minchin PR, et al. vegan: Community Ecology Package [Internet]. 2022. Available from: https://CRAN.R-project.org/package=vegan

