## Supplementary material for "Expansion, restructuring and characterization of the *Legionellaceae* family": Supplemetary information

**Supplementary text 1 – Comparison between proposed genus clusters and previous analyses**

Parks and coworkers (1) suggested to split the *Legionellaceae* family in 35 genera (as present in GTDB release 11-RS232) based on their relative evolutionary divergence. The even greater number of genera is due to the lower genome quality requirements required by GTDB compared to this study, as well as the reliance solely on RED values to define genera. Despite this, genus cluster boundaries are largely coherent between what proposed in this study and GTDB, showing an adjusted Rand Index of 0.87. Furthermore, the presence of multiple *Legionellaceae* clades was also reported in previous studies dedicated to these organisms (2). While such repartition into clades was not based on AAI nor POCP values, the proposed clades have a strong degree of concordance with the genus clusters identified by this work and differ only by the further separation of the genus clusters including *L. pneumophila* and *L. micdadei* (genus clusters 1 and 6) in two subclades. As all comparisons across subclades report POCP values above 50%, we believe that such subclades would result in an artificial overestimation of the number of *Legionellaceae* genera. In conclusion, the fact that comparable clades were identified across studies using different methodologies supports the robustness of our results and highlight their value for a detailed restructuring of this family in multiple genera.

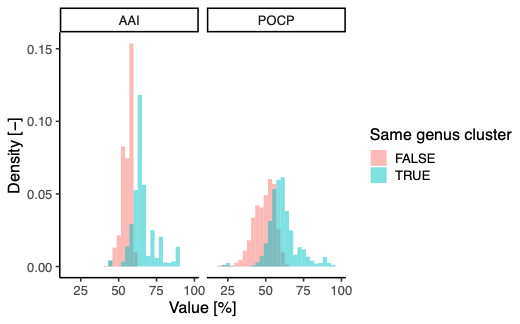

**Figure S1**. Distribution of AAI and POCP values across genus clusters. Values of POCP below 50% within the same genus cluster are due to the genome size reduction in *L. polyplacis*.

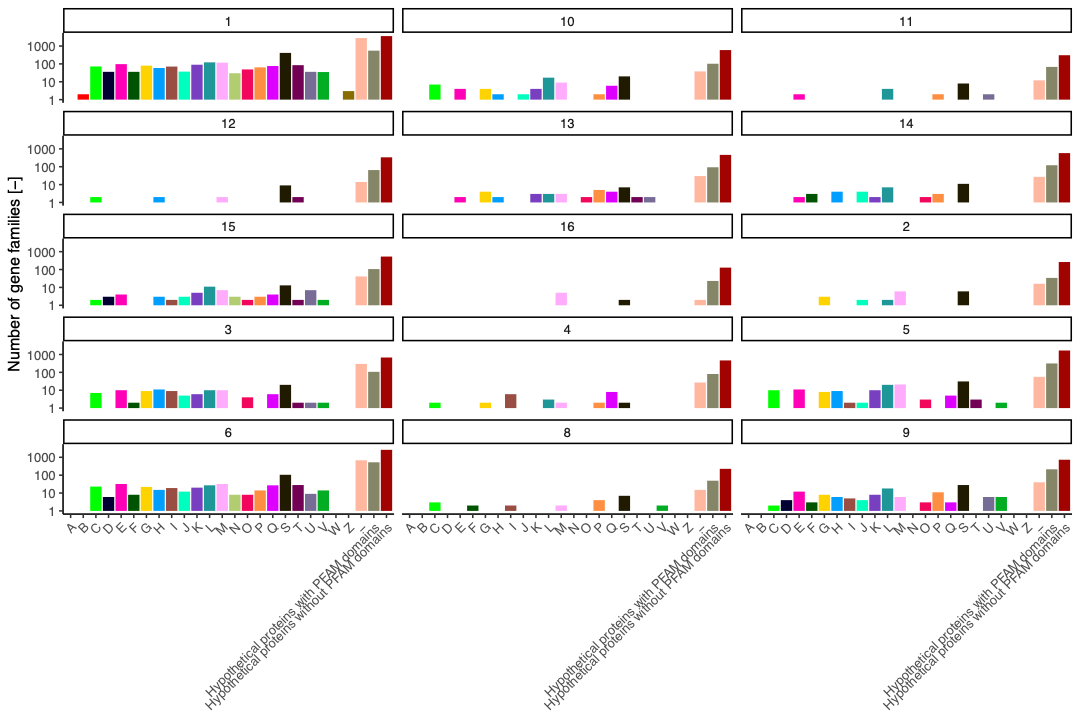
**Figure S2.** Number of novel protein families found uniquely in single genus clusters divided based on their COG class. Each facet refers to a single genus cluster.

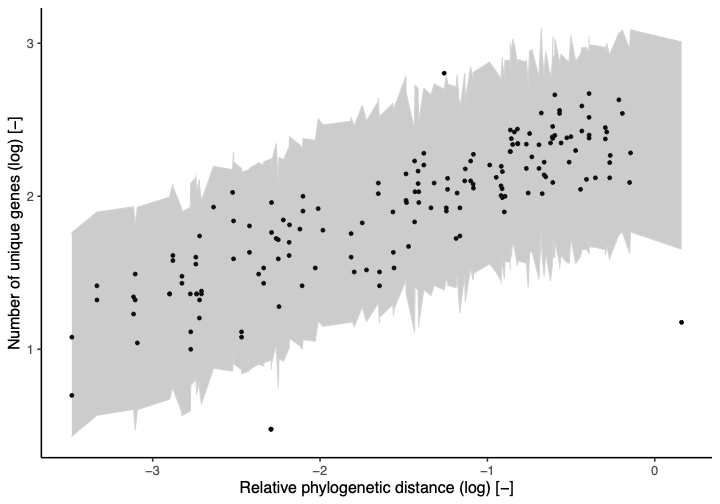

**Figure S3.** Number of unique genes per genome in relationship with the relative phylogenetic distance to the closest other genome analyzed. The area shaded in gray indicates the predicted 99% uncertainty interval.

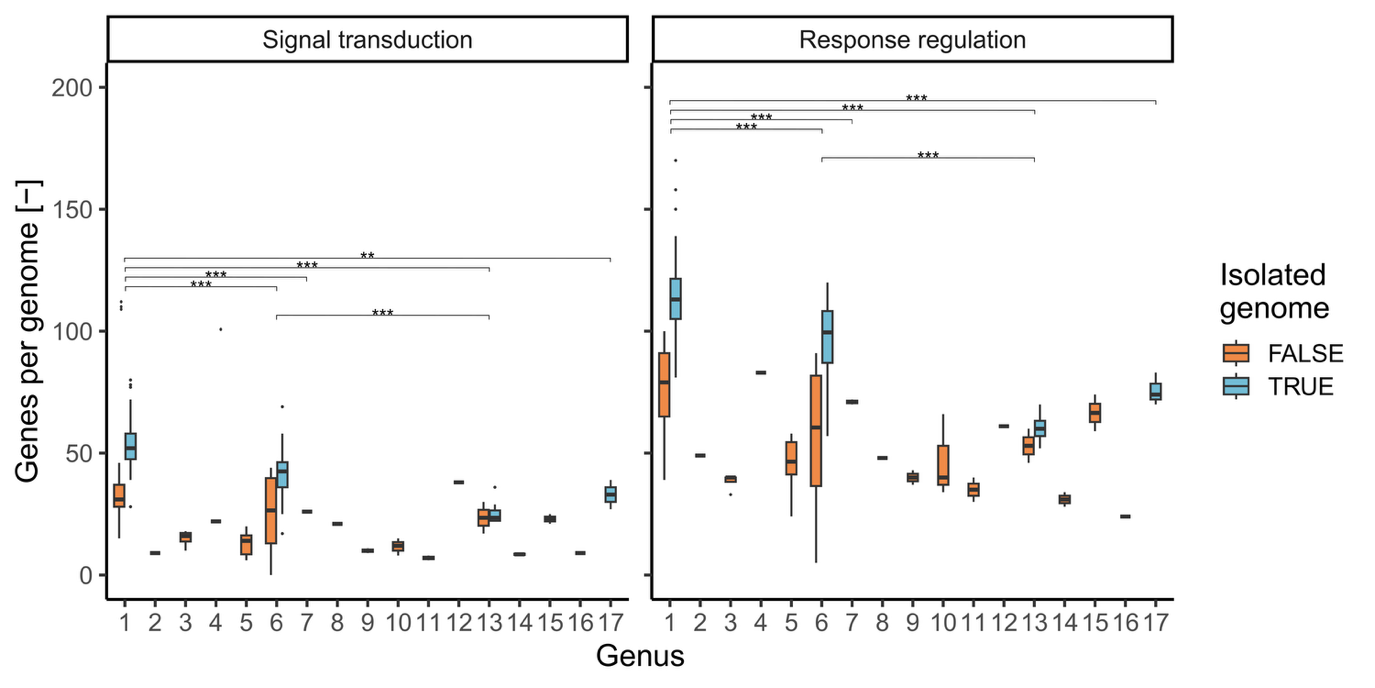
**Figure S4.** Number of signal transduction and response regulation detected per genome depending on genome source across different genera. Boxplot pairs marked with asterisks indicate statistically significant differences in the gene number detected in isolated genome from such genera (ANOVA, Tukey HSD post-hoc test).

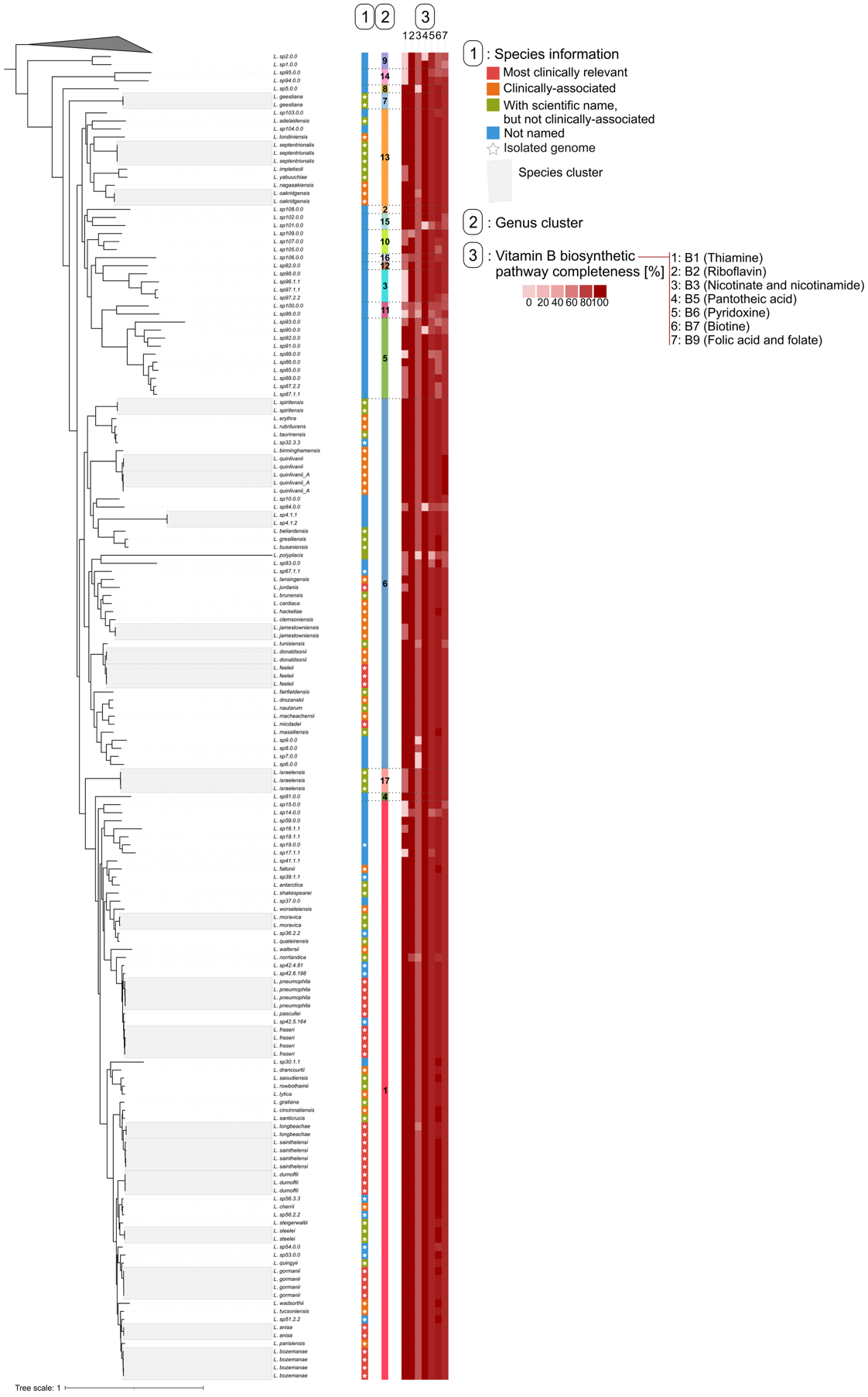

**Figure S5.** Maximum likelihood phylogenetic tree of the family *Legionellaceae* based on 369 BUSCO *Legionellales* protein families (123421 sites). Leaves enclosed within grey areas indicate genomes of different genomovars belonging to the same species, while shades of red indicate the completeness of vitamin B biosynthetic pathways.

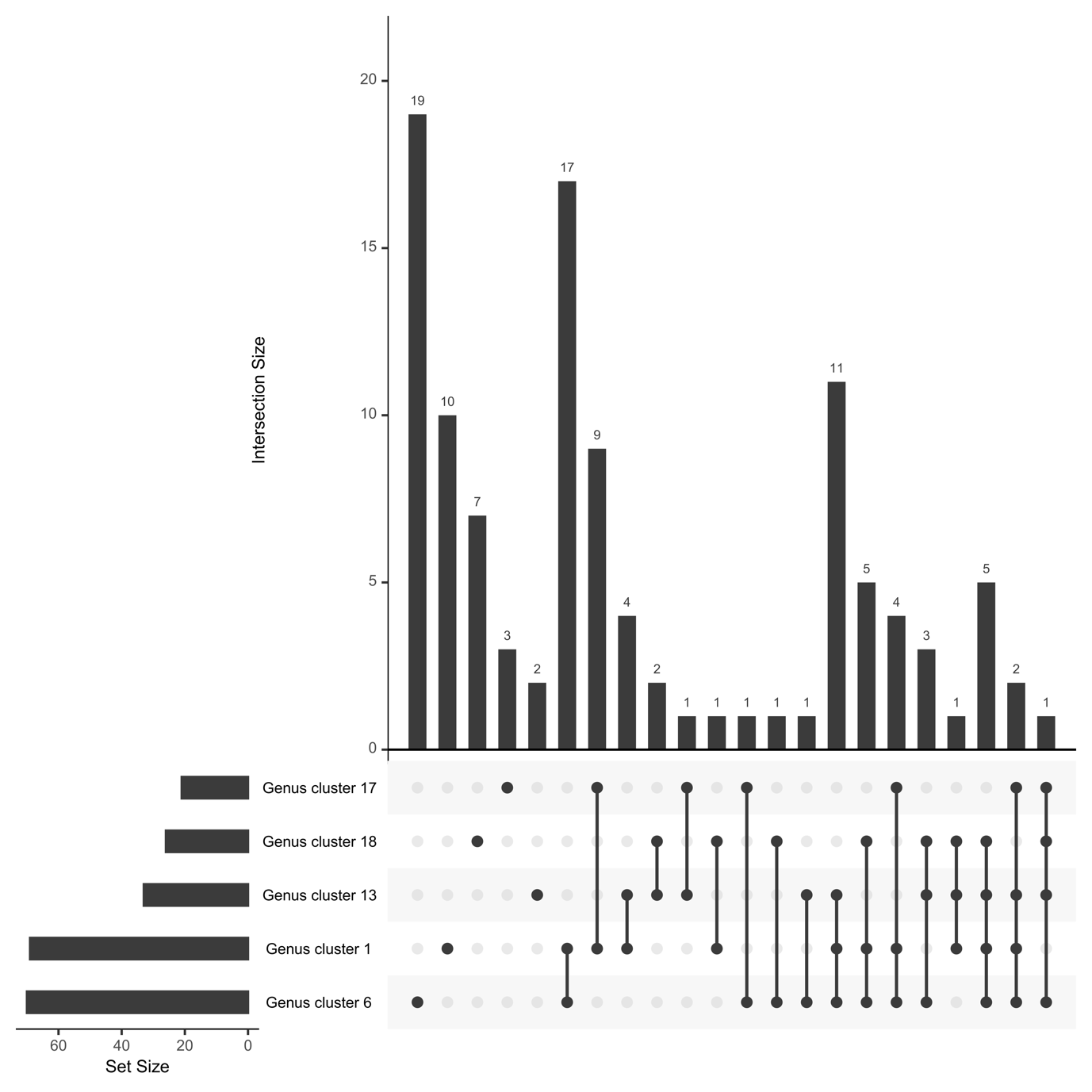
**Figure S6.** Intersection of the KOs which discriminate the genome content of each genus cluster against the remaining ones. To remove the bias linked to MAGs, this analysis was performed only using isolated genomes.

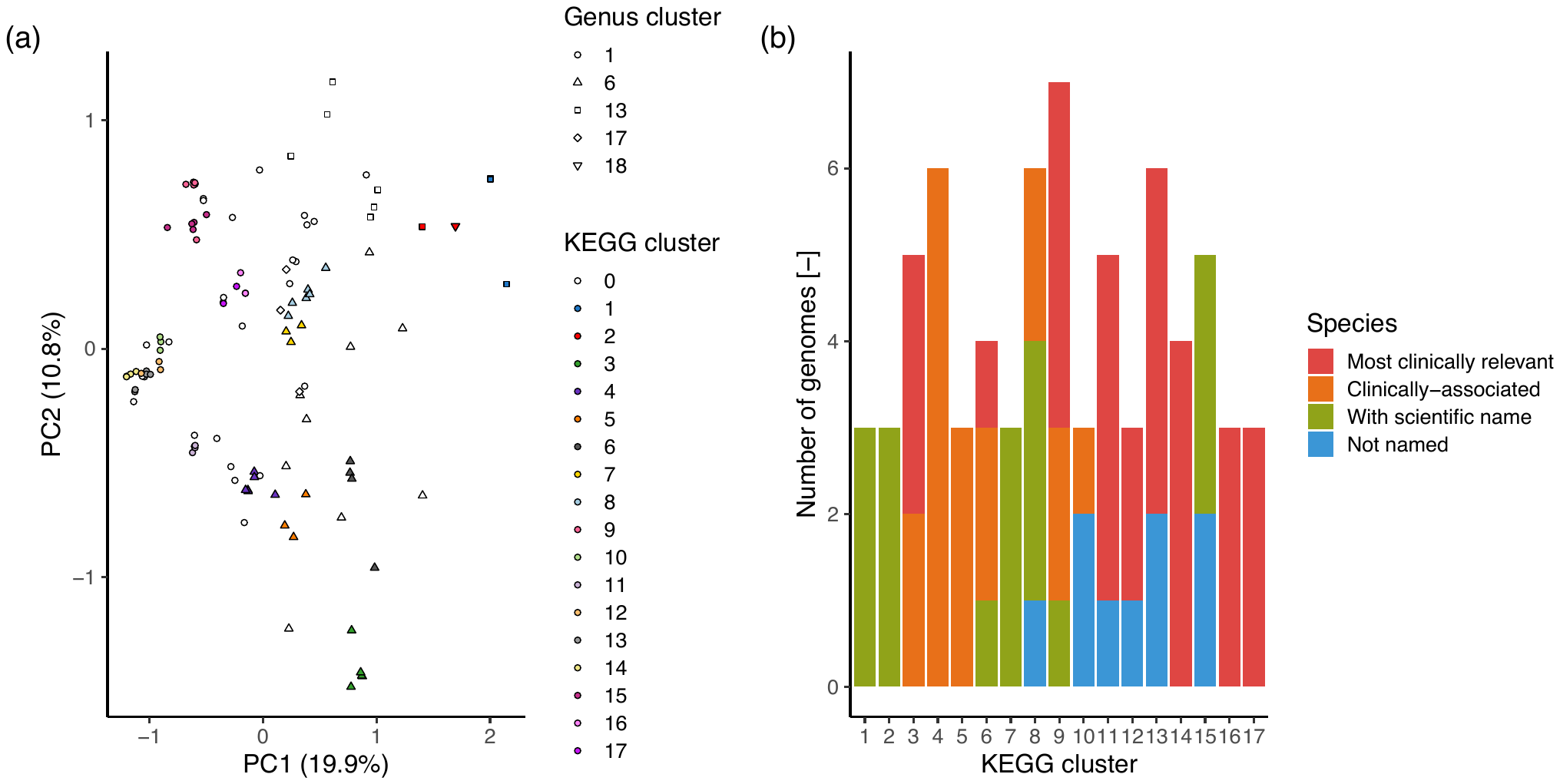
**Figure S7.** (a) Clusters of genomes with similar completeness of KEGG metabolic pathways. Cluster “0” indicates outliers, as defined by HDBSCAN. (b) Species information regarding the species belonging to each KEGG cluster.

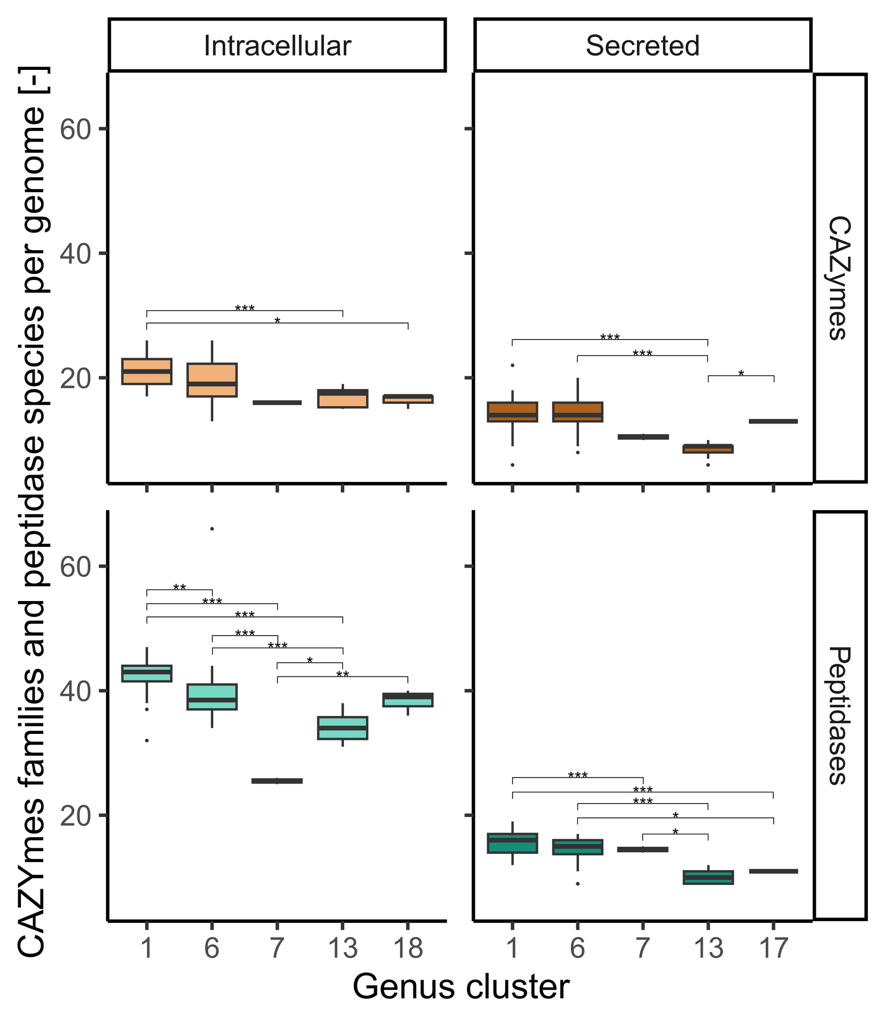

**Figure S8.** Number of CAZYmes families and peptidades classes in isolated genomes as a function of their secretion and genus cluster membership. Statistically significant differences among genus clusters are marked by their p-values: ”*” = 0.05 - 0.01, ”**” = 0.01 - 0.001, ”***” < 0.001.

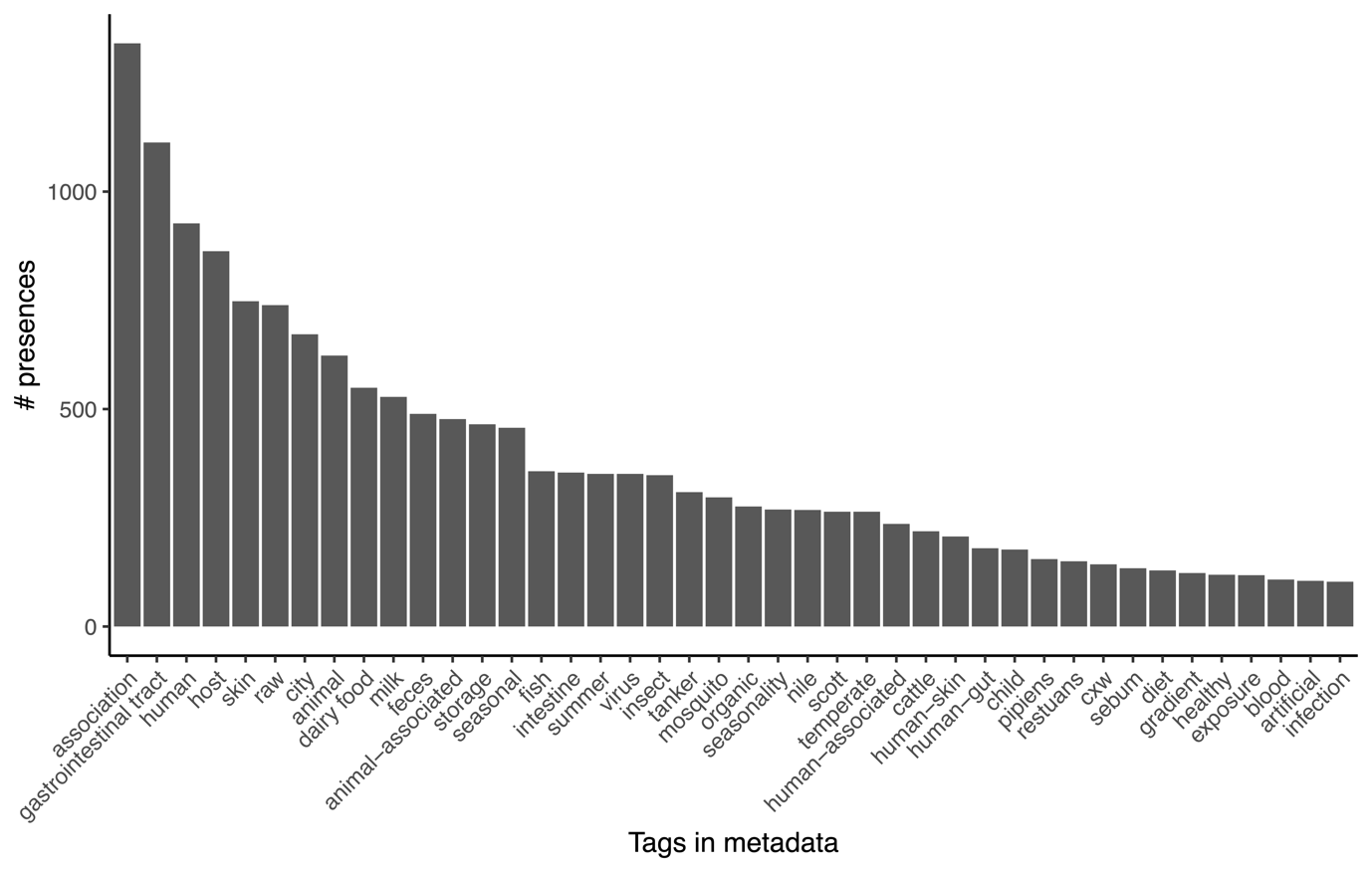
**Figure S9.** Number of appearances of the keywords linked to animal-associated samples containing the selected MicrobeAtlas OTUs.

References

1.	D. H. Parks, *et al.*, GTDB release 10: a complete and systematic taxonomy for 715 230 bacterial and 17 245 archaeal genomes. *Nucleic Acids Research* **54**, D743–D754 (2026).

2.	L. Gomez-Valero, *et al.*, More than 18,000 effectors in the *Legionella* genus genome provide multiple, independent combinations for replication in human cells. *Proc. Natl. Acad. Sci. U.S.A.* **116**, 2265–2273 (2019).
